# Non-specific adhesive forces between filaments and membraneless organelles

**DOI:** 10.1101/2021.07.22.453380

**Authors:** Thomas J. Böddeker, Kathryn A. Rosowski, Doris Berchtold, Leonidas Emmanouilidis, Yaning Han, Frédéric H. T. Allain, Robert W. Style, Lucas Pelkmans, Eric R. Dufresne

**Affiliations:** Department of Materials, ETH Zurich, Zurich, Switzerland; Department of Molecular Life Sciences, University of Zurich, Zurich, Switzerland; Institute of Biochemistry, ETH Zurich, Switzerland

## Abstract

Membraneless organelles are liquid-like domains that form inside living cells by phase-separation. While standard physical models of their formation assume their surroundings to be a simple liquid, the cytoplasm is an active viscoelastic environment. To investigate potential coupling of phase separation with the cytoskeleton, we quantify structural correlations of stress granules and microtubules in a human-derived epithelial cell line. We find that microtubule networks are significantly perturbed in the vicinity of stress granules, and that large stress granules conform to the local pore-structure of the microtubule network. When microtubules are depolymerized by nocodazole, tubulin enrichment is localized near the surface of stress granules. We interpret these data using a thermodynamic model of partitioning of particles to the surface and bulk of droplets. This analysis shows that proteins generically have a non-specific affinity for droplet interfaces, which becomes most apparent when they weakly partition to the bulk of droplets and have a large molecular weight. In this framework, our data is consistent with a weak (≲ *k_b_T*) affinity of tubulin sub-units for stress granule interfaces. As microtubules polymerize their affinity for interfaces increases, providing sufficient adhesion to deform droplets and/or the network. We validate this basic physical phenomena *in vitro* through the interaction of a simple protein-RNA condensate with tubulin and microtubules.

---

Phase separation is a physical mechanism utilized by cells to rapidly alter the biochemical landscape by locally concentrating or segregating key molecules [1, 2]. The resulting domains are called membrane-less organelles and are involved in various processes inside the cell. Typically composed of protein and RNA, they include, for example, nucleoli in the nucleus as well as p-bodies and stress granules in the cytoplasm [3, 4]. Membrane-less organelles can be very dynamic, exhibiting liquid properties such as coalescence [5–8] and recovery from photobleaching [4, 5, 9]. In recent years, biologists, chemists and physicists have come together to understand the interplay of phase separation, composition and function of these droplet-like domains [10].

Phase separation inside cells takes place in an active, viscoelastic environment. Be it the chromatin network of the nucleus or the cytoskeleton of the cytoplasm, protein-mRNA droplets grow within filamentous networks [11]. Recent studies have focused on the impact of networks on the growth of droplets at scales well beyond the mesh scale. In that case, elastic energy stored in network deformations has been found to significantly alter droplet growth and coarsening, both for synthetic mixtures of oil in cross-linked silicone [12–14] and protein droplets in chromatin [15–18].

In the cytoplasm, membrane-less organelles interact with filaments of the cytoskeleton. For example, multiple lines of cell-biological evidence show that stress granules can interact with microtubules directly or through interaction with microtubule-binding p-bodies [5, 19]. Stress granules are liquid-like complexes of proteins and mRNA [20] that form throughout the cytosol under conditions of biological stress (*e.g.* heat shock or exposure to arsenite), that modulate the translation of cytoplasmic mRNA [4, 9, 21, 22]. As biological stress persists, these granules grow and coalesce to reach a size up to a few microns [23, 24]. Microtubules have been suggested to aid stress granule formation by acting as tracks for active transport of granule components [25–28] and by encouraging droplet fusion [23, 24, 29]. Since the characteristic mesh size of the microtubule network is about a micron, the dimension of membraneless organelles and the microtubule mesh are comparable, and their interactions fall in an unexplored physical regime. Recent theoretical works suggest that structure and mechanical properties at the pore-scale can play an essential role [30, 31].

Here, we investigate physical mechanisms underlying the interaction of microtubules and stress granules. Analysis of their structural correlations shows that microtubule network densities are enhanced at the surface of stress granules and decay to their usual value over distances much larger than the granule size. Conversely, large stress granules tend to conform to irregular gaps in the network of microtubules. When microtubules are perturbed by nocadozole, which hinders microtubule polymerization, tubulin intensities remain enhanced, but more strongly localized, at the surface of stress granules. These observations point to adhesive interactions of tubulin and stress granules. A simple and generic model shows that surface adsorption requires no surface-specific molecular functionalization. Instead, it emerges naturally when macromolecular sub-units, with little preference for either phase, assemble into larger structures. While these non-specific adsorption energies are very small (≲ 0.1*k_b_T*) for a single tubulin sub-unit, they are sufficient to cause adhesion between microtubules and the surface of stress granules with a strength of order 50 *k_B_T/μ*m. We demonstrate this effect *in vitro* by comparing the interaction of RNA-protein condensates with tubulin sub-units or microtubules.

To reveal interactions of membrane-less organelles and cytoskeletal filaments, we chose to work with stress granules in U2OS epithelial cells. On fibronectin functionalized substrates, U2OS spread thoroughly, facilitating cytoskeletal imaging [32]. Formation of stress granules is readily triggered by addition of small quantities of arsenite to the imaging media [5, 21]. An example of the nucleation and growth of stress granules in a live cell tagged for G3BP1 (a stress granule marker protein) and tubulin is shown Supplementary Movie 1 (see supplement section A). This movie suggests a rich interplay of microtubules and stress granules, which not only varies over time as the granules grow, but also across different positions in the cell. To quantify these interactions we record confocal stacks of fixed cells after 90 minutes of arsenite treatment. We find that granules form within about 15 to 30 minutes after treatment, continue to coalesce and grow but show little change in size after about 60 minutes of treatment. To limit spatial variation, we controlled cell shapes by plating them on patterned fibronectin patches [32]. All cells whose final shape did not conform to the pattern were excluded from analysis. We stained cells for G3BP1 and *β*-tubulin and acquired stacks of *xy* images separated by Δ*z* = 0.2 *μ*m with a spinning disc confocal microscope (Nikon Ti2 with Yokogawa CSU-W1, 100x, NA 1.45).

An example *xy*-slice of the G3BP1 and *β*-tubulin channels 90 minutes after arsenite treatment are shown in Figure 1 (*A,B*). In this example, we notice several important features. Stress granules have a wide range of sizes, with compact irregular shapes. They tend to be found in the region near the nucleus, which is also rich in microtubules. To quantify these observations, we automatically detect individual stress granules and record their spatial coordinates, shape, and size (see supplement section B5). We find a size distribution of granule radii with a broad peak between 0.5 and 1 μm, which decays towards larger sizes with practically no granules larger than 1.5 μm (see Fig. S7). We classify granule shape as round or elongated. Round granules have an aspect ratio between 1 to 1.5, and make up 79% of observed stress granules. Elongated granules have aspect ratios greater than 1.5, and can reach values as large as 5.2 (Fig. S7). Live cell imaging of GFP-G3BP1 tagged stress granules confirms that these large aspect ratios can relax over time, and are therefore not a sign of granule hardening (Supplemental Movie 1).

**FIG. 1.**
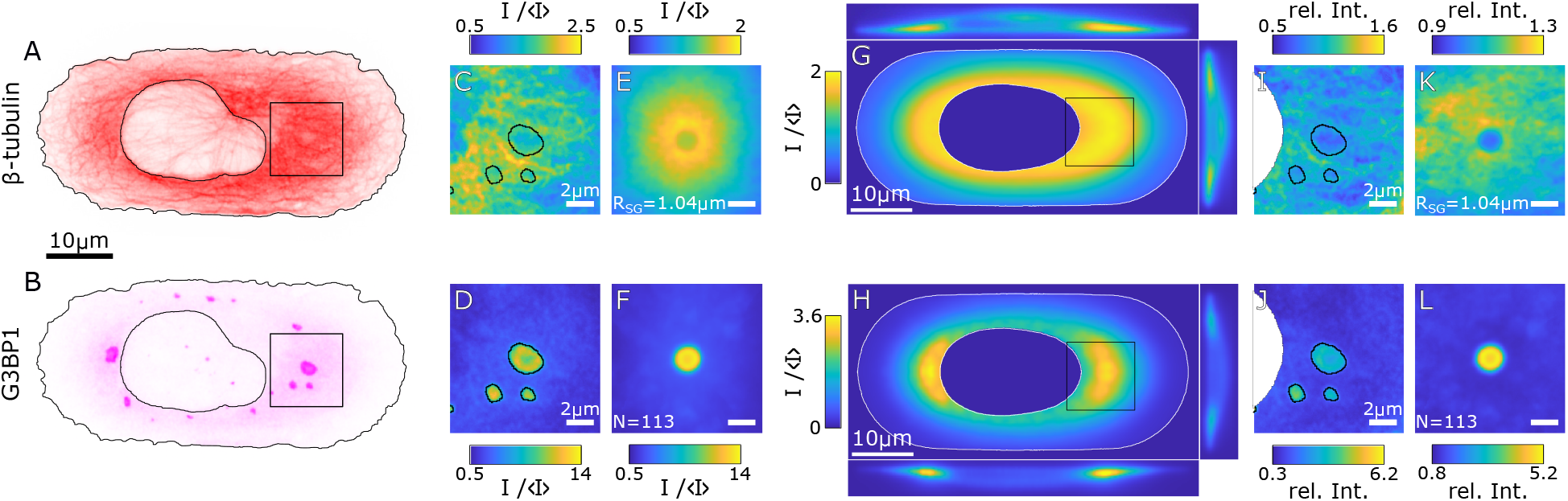
Stress granules and the surrounding microtubule network. (*A,B*) Exemplary xy-slices of the G3BP1 and *β*-tubulin channels of the same cell after 90 minutes of arsenite treatment. The black line shows the outline of the detected cell and nucleus. (*C,D*) Images of the stress granule and surrounding microtubule network at the location indicated by the box in (*A,B*). 〈*I*〉 gives the average intensity of the respective channel across a cell (see supplement section B6). (*E,F*) Averaged images of *N* = 113 stress granules and their microtubule network for granules with a radius within the interval of 0.98 to 1.11 μm. (*G,H*) Exemplary xy-slices of the reference cells calculated from *N* = 335 cells for the G3BP1 and *β*-tubulin channel as well as side views along the respective center line. The white line indicates the average cell and nucleus shape. The z-axis has been scaled to match the resolution in the xy-plane. (*I,J*) Same images as in (*C,D*) normalized by the intensities of the corresponding location in the respective reference cell (indicated by the black boxes). The intensity is now given in units of *relative intensity*, where a value of one corresponds to *as expected in the reference cell. (K,L)* Distribution maps of G3BP1 and *β*-tubulin for granules with a radius within the interval of 0.98 to 1.11 μm.

To reveal interactions of stress granules and microtubules, we quantify the structure of the microtubule network in the presence of stress granules. The basis of this analysis are pairs of images of the G3BP1 and *β*-tubulin channels centered on a given granule. An example of such a image pair is shown in Fig. 1 (*C*) and (*D*). For the moment, we consider only round granules with radii greater than 195 nm. This corresponds to the diffraction limit of our optical setup (see supplement section B4). To visualize the typical shape of both granules and their surrounding network, we average the granule-centered images of both channels for granules of similar radii. Example average images for granules with effective radii of about 1.04 μm are shown in Fig. 1 (*E*) and (*F*). The average images for all size bins can be found in Fig. S3 and S4.

The average image of the stress granule channel shows a circular granule with an intensity greater than 10 × the average G3BP1 intensity in the cell. The average *β*-tubulin images reveal a volcano-like shape, with a central region of relatively lower intensity that matches the size and shape of the granule. This circular region is surrounded by a ring-like structure of almost twice the normal tubulin concentration, followed by a decay in *β*-tubulin intensity over several micrometers towards the average tubulin intensity.

To quantify how local perturbations of microtubule network structure are correlated with the presence of stress granules, we needed to account for systematic variations in microtubule architecture across the cell. To do so, we constructed *reference cells*, spatially-resolved image stacks for each channel averaged across the full ensemble of cells. Details of image alignment, construction and normalization are described in supplement section B8. The reference cells provide the expected intensity of the *β*-tubulin and G3BP1 channels for each 3D location in the cell, as well as the average cell shape and location of the nucleus. Slices of the G3BP1 and *β*-tubulin reference cells in the *xy, xz* and *yz*-planes are shown in Fig. 1 (*G*) and (*H*). Both tubulin and G3BP1 are brightest near the nucleus.

With reference cells in place, we calculate the enhancement of G3BP1 and *β*-tubulin intensity in and around individual granules by point-wise division of the stress granule or *β*-tubulin images by the intensity of the corresponding reference cell at the same location. Figure 1 (*I*) and (*J*) show the same images as 1 (*C*) and (*D*), where each pixel is now normalized by the respective reference. Thus, a value greater (less) than one indicates that the intensity is higher (lower) than expected at that location. Note that pixels that are masked due to being outside the cell or inside the nucleus in either individual image and/or the reference are discarded.

Averaging these normalized images for the same granule size and condition, we arrive at *distribution maps*, see Fig. 1 (*K*) and (*L*). Qualitatively, the average images (Fig. 1 (*E*)) and distribution maps for the G3BP1 channel are quite similar. The normalization largely just rescales the relative intensities. For the tubulin channel, normalization by the local intensity has stronger effect. It flattens the intensity distribution outside of the stress granule, while retaining the enhancement of tubulin around its surface. Within the granule, the tubulin signal drops to a value close to one.

These basic features are found for granules of all sizes, as shown in Fig. 2. However, the enhancement of the microtubule signal around stress granules is stronger for larger granules, as exemplified in Fig. 2 (*A-D*). For quantitative comparison, we azimuthally average the distribution maps, to form a radial distribution function *g*(*r*) for the tubulin and stress granule channels. These curves are shown for various granule radii in Fig. 2 (*E*) and (*F*). As expected for a compact phase-separated object, *g*(*r*) for G3BP1 peaks near *r* = 0 and decays to one over a distance corresponding to the granule radius. For *β*-tubulin, *g*(*r*) increases from a value near one at the center of the granule (*r* = 0) to a peak just outside the granule, followed by a slow monotonic decay over a distance much larger than the size of the granule.

**FIG. 2.**
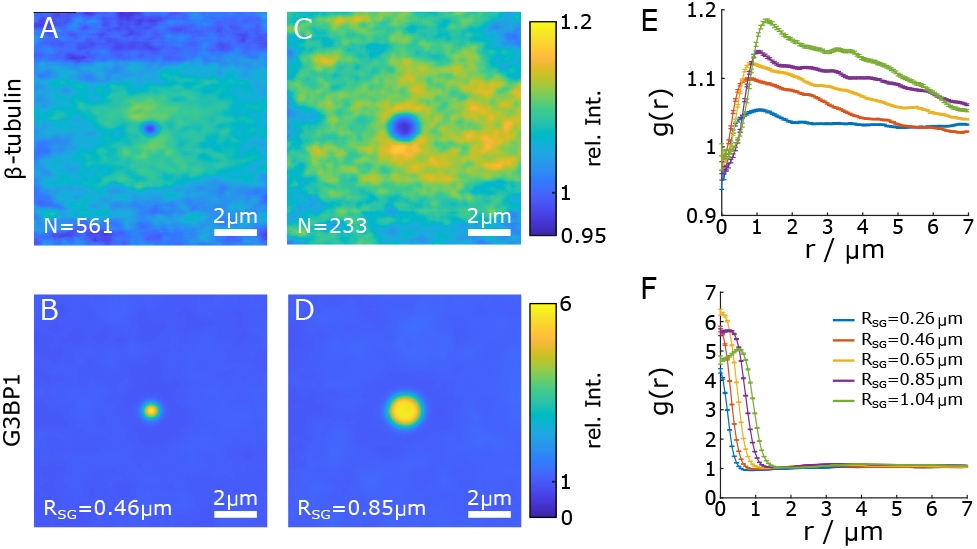
The microtubule network is modulated in the presence of stress granules. (*A*) distribution map of the *β*-tubulin channel for round granules with a radius from 0.39 to 0.52 μm. *N* gives the number of contributing images. (*B*) Corresponding distribution map for G3BP1. (*C*) distribution map of the *β*-tubulin channel for round granules with a radius from 0.78 to 0.91 μm. (*D*) Corresponding distribution map for G3BP1. (*E*) Radial distribution function *g*(*r*) of *β*-tubulin around round granules of different size. (*F*) *g*(*r*) of G3BP1.

We validated this novel method to characterize structural correlations in a heterogeneous environment with a computational negative control. In order to construct distribution maps with no structural correlations, we used the measured positions of each stress granule, but images of the G3BP1 and tubulin channels at the same location in a different cell. As expected, the resulting distribution maps for both G3BP1 and *β*-tubulin showed none of the features described above, but simply fluctuate around a value of one, see Fig. S3 and S4.

So far, we have considered only granules with a principal axis ratio up to 1.5. Now, we turn our attention to the network around elongated granules. Figure 3 shows the distribution maps for granules with larger ellipticity. Here, we keep the size of the granule constant and bin the granules by their principal axis ratio. The construction is analogous to the distribution maps of round granules, only now individual images of granules, as well as the corresponding image of the tubulin channel, are rotated such that the major principal axis is vertical before averaging. We find that elongated granules are correlated with complementary depletions in the *β*-tubulin intensity that match its size and shape. On average, larger stress granule are more elongated, suggesting that larger granules are more affected by the network, see Fig. S7 (*H*). Since the favored shape of a liquid droplet is spherical, elongated droplets reflect anisotropic forces acting upon them. The correspondence in shape of the granules and the ‘cavity’ in the tubulin channel suggests that the microtubule networks can deform stress granules.

**FIG. 3.**
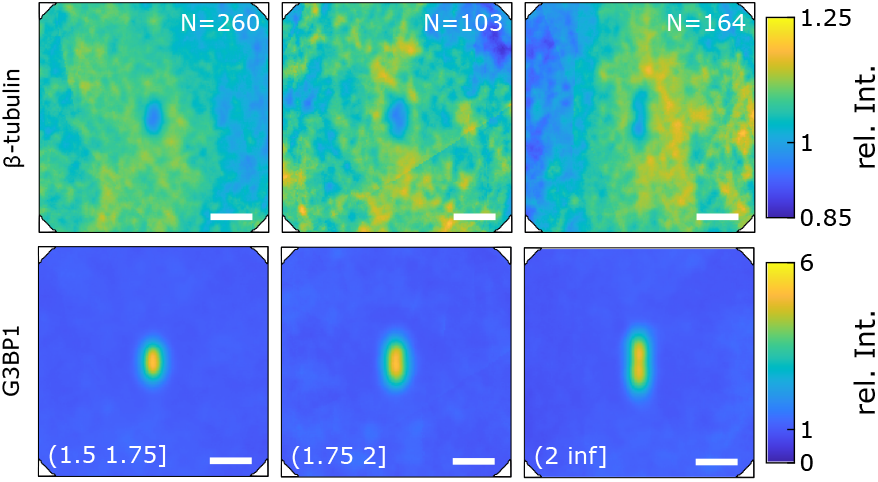
Elliptical granules correlate with elliptical shapes of the microtubule network. Distribution map of the tubulin network and corresponding granules around granules binned by principal axis ratio. The area of all granules is fixed between 0.5 and 1.5 μm^2^. Both individual granule as well as tubulin image are rotated such that the long axis of each granule is vertical.

Our results suggest that microtubule networks are deformed in the presence of stress granules (Fig. 2) and that stress granules conform to the local structure of the microtubule network (Fig. 3). To perturb microtubules, we treat cells with 0.5 μg/ml nocodazole. Nocodazole hinders microtubule polymerization, leading to depolymerization of the network within 10 to 20 minutes [33]. Cells are treated for 90 minutes with both sodium arsenic and nocodazole before fixation. After nocodazole treatment, filamentous structures are absent from the *β*-tubulin channel and stress granules are smaller and more numerous (Fig. S7), in line with previous observations [23]. Note that we can not confirm that microtubules are fully depolymerized into sub-units, tubulin may also be present in the form of small aggregates below the optical limit.

Distribution maps of nocodazole-treated cells are shown in Fig. 4 (*A, B*). The distribution map of *β*-tubulin shows a relatively narrow ring surrounding the peak in the G3BP1 channel. This observation becomes clearer in *g*(*r*), which reveals a peak localized at the granule surface. A direct comparison of the *β*-tubulin and G3BP1 *g*(*r*)’s (Fig. 4 (*C*)) reveals that the surface enhancement of *β*-tubulin is more localized in nocodazole-treated cells. Further, the maximum value of the *β*-tubulin *g*(*r*) is typically higher in nocodazole-treated cells for granules of all sizes (Fig. 4 (*D*)). Finally, the location of the *β*-tubulin peak, *r*_max_ is consistently closer to the center of the granule in nocodazole-treated cells (Fig. 4 (*E*)).

**FIG. 4.**
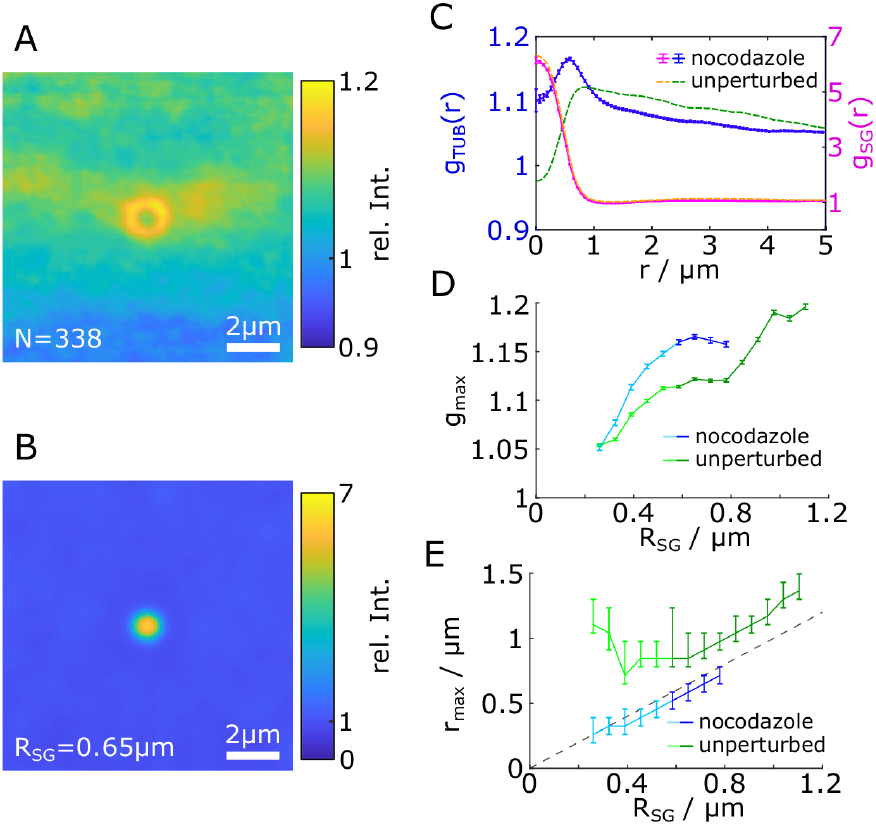
*β*-tubulin localizes at the granule surface in nocodazole-treated cells. Distribution map of *β*-tubulin (*A*) and G3BP1 (*B*) of granules with a radius around 0.65 μm in nocodazole-treated cells. (*C*) *g*(*r*) for the distribution maps in the first two panels (solid line with error bars) compared to the *g*(*r*) of the same size of granules with intact microtubule network (dashed lines). (*D*) Maximum value, *g*_max_, of the tubulin *g*(*r*) for granules of different radius calculated as the running average considering data of the indicated radius ±0.065 μm. Data are shown for both nocodazole-treated cells and cells with arsenite treatment only. The darker color shades indicate the size of granules where the intensity of G3BP1 saturates inside the granule. (*E*) Running average of the position of the maximum value *r*_max_ in the tubulin *g*(*r*) for granules of different radius in nocodazole-treated and untreated cells. The dashed line indicates the detected granule surface.

The persistent preference of tubulin, with or without a well-defined microtubule network, for the surface of stress granules suggests that tubulin may have an affinity for the surface of stress granules. To investigate this more precisely, we quantify intensities as a function of distance, *d*, from the stress granule surface, instead of the distance, *r*, from the center of the granule. In this way, we can consolidate data from granules of different sizes and shapes. The interface (*d* = 0) is defined by the granule detection routine. Averaging intensities for pixels with the same *d* from reference-cell-normalized images, we arrive at a surface-relative distribution function, *g_s_*(*d*), for each granule. An example of *g_s_*(*d*) for the G3BP1 and *β*-tubulin channels of one granule in a nocodazole-treated cell is shown in Fig. 5 (*A*). *g_s_*(*d*) for the G3BP1 channel reveals an interface region over which the intensity of G3BP1 transitions smoothly from the inside of the granule to the cytosol. The corresponding *g_s_*(*d*) for tubulin has a strong peak within this interface zone. While there is strong variation in the *β*-tubulin *g_s_*(*d*) from granule to granule (Fig. 5 (*B*)), we typically observe a local maximum within this interface zone. This observation becomes clear in the average *g_s_*(*d*) of *β*-tubulin, which shows a peak centered within the interface zone of the G3BP1 *g_s_*(*d*) (Fig. 5 (*C*)).

**FIG. 5.**
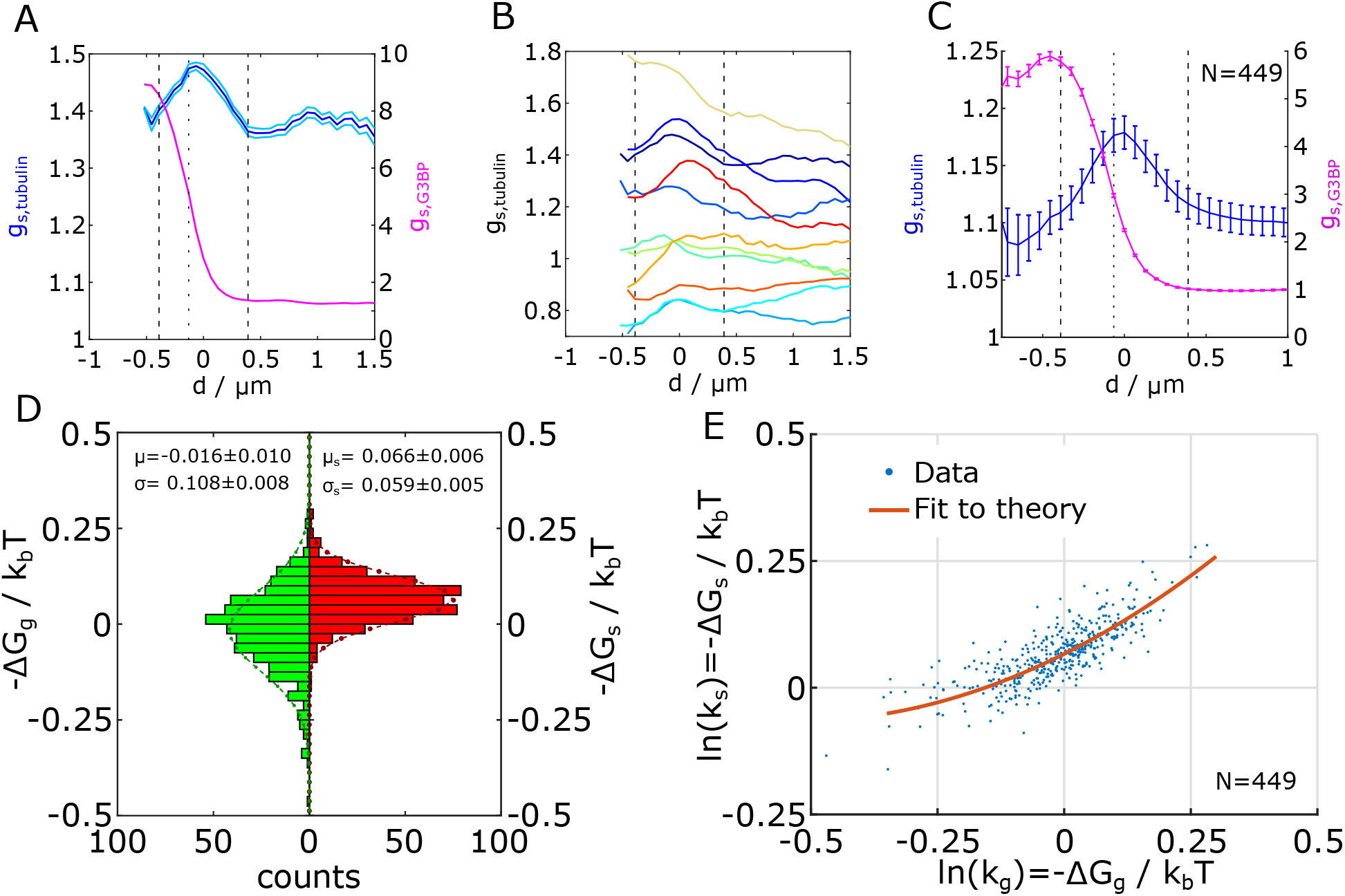
Tubulin sub-units in nocodazole-treated cells adhere to the granule surface. (*A*) Surface-relative distribution function *g_s_*(*d*) for the *β*-tubulin as well as the G3BP1 channel for a single stress granule in a cell treated with nocodazole. The dashed vertical lines at ±0.39 μm indicate the approximated width of the surface enhancement. The dotted vertical line shows the position of the maximal slope in the G3BP1 channel. (*B*) *g_s_*(*d*) for the *β*-tubulin for randomly picked granules. (*C*) *g_s_*(*d*) of the tubulin and G3BP1 channels for all sufficiently large granules (*N* = 449), i.e. collected over all data shown in the scatter plot in panel (*E*). The dashed lines show the assumed width of the interface at ±0.39 μm and the dotted line shows the position of the maximal gradient in *g_s,G3BP_*(*d*). (*D*) Histogram of the natural logarithm of the bulk partitioning coefficient ln(*k_g_*) = – Δ*G_g_* and the surface partitioning ln(*k_s_*) = –Δ*G_s_* with the respective mean and variance of a Gaussian fit. (*E*) Scatter of ln(*k_s_*) as a function of ln(*k_g_*). Each point corresponds to one granule (*N* = 449). The red line shows the best fit of the theory to the data.

We quantified the affinities of tubulin sub-units for the surface and bulk of stress granules using partition coefficients. Generally speaking, partition coefficients measure the concentration of a species in a particular domain relative to their values in a nearby reservoir, here the cytosol. Based on the average *g_s_*(*d*) for G3BP1, we define the interface zone of width |*d*| < 0.39 *μ*m, where G3BP1 intensities transition from their values in the stress granule to their values in the bulk. We defined the surface partition coefficient, *k_s_*, as the ratio of the peak value of *g_s_*(*d*) within the interface zone to its value just beyond the interface zone, outside of the granule. Similarly, the partition coefficient for the bulk of an individual granule, *k_g_*, is the ratio of *g_s_*(*d*) just inside to just outside the interface zone (see supplement section B10). Given the broad apparent interface region, we only consider granules with a radius of 0.59 μm and larger, i.e. granules with a defined bulk phase. Note that these partition coefficients are based on intensities, not concentrations. Therefore, systematic variations in fluorophore efficiency or antibody binding in different environments could lead to systematic errors in partition coefficients. However, we expect such errors to be negligible in the limit of weak bulk partitioning (*k_g_* ≈ 1) as observed here.

Assuming local equilibrium, we can relate the partition coefficients to the free energy difference of tubulin relative to the cytosol:

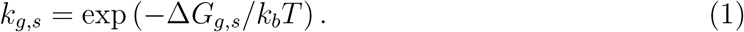

Here, Δ*G_g_* and Δ*G_s_* indicate the free energy difference of tubulin inside and on the surface of stress granules, respectively, relative to the cytosol. Histograms of the bulk and surface affinities of tubulin — Δ*G_g_* and — Δ*G_s_* are shown in Fig. 5 (*D*). We find that the distribution of affinities for the bulk of the granules has a small but statistically significant (*p* = 0.002) mean of —0.016 *k_b_T*, suggesting a subtle preference for the cytosol. On the other hand, the distribution of surface affinities has a significant (*p* < 0.001) mean value of 0.066 *k_b_T*. This suggests that tubulin sub-units have a weak attraction to the interface of stress granules. The apparent surface and bulk affinities are correlated, as shown by the scatter plot in Fig. 5 (*E*). The surface affinity increases with the bulk affinity, but remains positive as the bulk affinity vanishes.

This behavior is consistent with a simple physical picture where tubulin is attracted to the surface to reduce the interfacial energy of stress granules. In other words, tubulin acts as a very weak surfactant. To capture the essential physics, we developed minimal physical models where tubulin (in the form of a sub-unit or a small aggregate) has no specific molecular mechanism for attachment to the surface of a stress granule. As in the theory of Pickering emulsions [34–36], ûbulin’s affinity for the bulk of the granule is Δ*G_g_* = *A*_0_Δ*γ* = *A*_0_(*γ_tg_* — *γ_tc_*). Here, *A*_0_ is the surface area of the tubulin, and *γ_tg_* & *γ_tc_* are the interfacial energies of the tubulin against the granule & cytoplasm, respectiviely. In the special case where the tubulin would have no preference for either phase (—Δ*G_g_* = 0), it would be bound to the interface with an affinity — Δ*G_s_* = *A_x_γ_cg_*, where *A_x_* is the area of the interface covered by the tubulin sub-unit and *γ_cg_* is the interfacial energy between both liquid phases (see supplement section D). The affinity is simply caused by the reduction of energy of the interface when part of the interface is covered by a particle. When the particle is larger, it takes up more area at the interface and is stronger bound. Assuming that tubulin attaches to the interface in sub-units with *A_cut_* ≈ 28 nm^2^ [37], the observed surface affinity at Δ*G_g_* = 0 of about 0.1 *k_b_T* is consistent with a interfacial energy of the stress granules *γ_cg_* ≈ 15 *μ*J/m^2^, which is well within the range of typical values for membraneless organelles [6, 38]. Since a single sub-unit is the smallest plausible aggregate size, this estimate is an upper bound for the interfacial energy.

The theory of Pickering emulsions, however, was developed for systems where the particle size is much larger than the interface thickness. Since tubulin sub-units have a similar size to the components of stress granules, we do not expect it to apply here. We further note that the apparent half-width of the interface is about 0.4 μm (Fig. 5 (*C*)), which is much larger than the size of a tubulin sub-unit. Therefore, we extend the basic Pickering concept to situations where particles interact with an interface of finite width, *w*. In the limit where the particle size is much smaller than *w*, the free energy difference between a particle at position *x* and bulk cytosol (*x* → ∞) has two terms, one proportional to its surface area, *A*_0_, and one to its volume, *V*_0_:

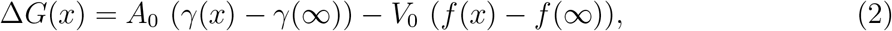

The area term captures the interfacial energy between particle and fluid and the volume term describes the energy of the fluid displaced by the particle. To determine the free energy of the particle localized to a thick interface, we follow the description of Cahn and Hillard of a simple two-component system [39–41]. Expanding the energy landscape around the center of the interface (*x* = 0) and keeping terms up to second order in *x/w*, we find

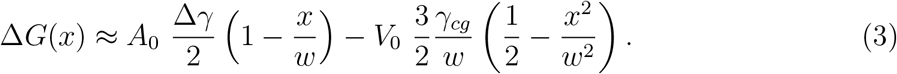

A particle at the interface then has the equilibrium position

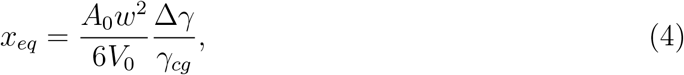

with a corresponding affinity

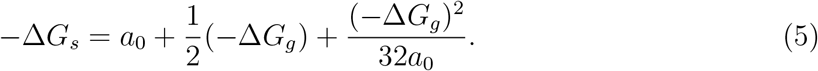

We find a single physical parameter *a*_0_, which gives the surface affinity of a tubulin particle at Δ*G_g_* = 0. For a spherical particle with radius *R*,

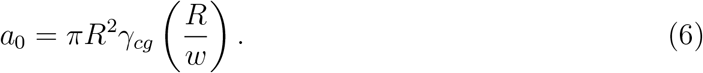

Note that the thin interface model gives a similar quadratic form with different pre-factors. Complete derivations of each theory are provided in supplement sections D and E. Fitting the data in Fig. 5 (*E*) to equation 5 with only one fit parameter, we find *a*_0_ = 0.067 ± 0.002 *k_b_T*, see supplement section B11.

The above analysis of nocodazole-treated cells shows that tubulin fragments have a small affinity (≲ 0.1 *k_b_T*) for the surface of stress granules. As tubulin polymerizes to form microtubules, this affinity should increase linearly with the length of the filament. Assuming that the surface affinity measured in nocodazole corresponds to individual sub-units, microtubules as short as 100nm will bind to the surface of stress granules with an adhesion energy of more than 5 *k_b_T*. This implies that microtubules can strongly bind to the surface of stress granules, even when tubulin sub-units do not.

To test this, we performed experiments on a simplified system *in vitro*. In place of stress granules, we made protein-RNA condensates with full length FUS protein, a component of stress granules [9], and RNA (poly-U), see supplement section B13. In this minimal system, we observed a significant affinity of rhodamine-labelled tubulin sub-units for the surface of the droplets, see Fig. 6 (*A-D*). Strikingly, taxol-stabilized microtubules have a much stronger preference for the surface of the droplet, see Fig. 6 (*E-I*). This is consistent with the simple model we have proposed, and is reminiscent of other recent *in vitro* observations, where filaments were strongly localized to the surface of synthetic phase-separated polymer droplets [42, 43].

**FIG. 6.**
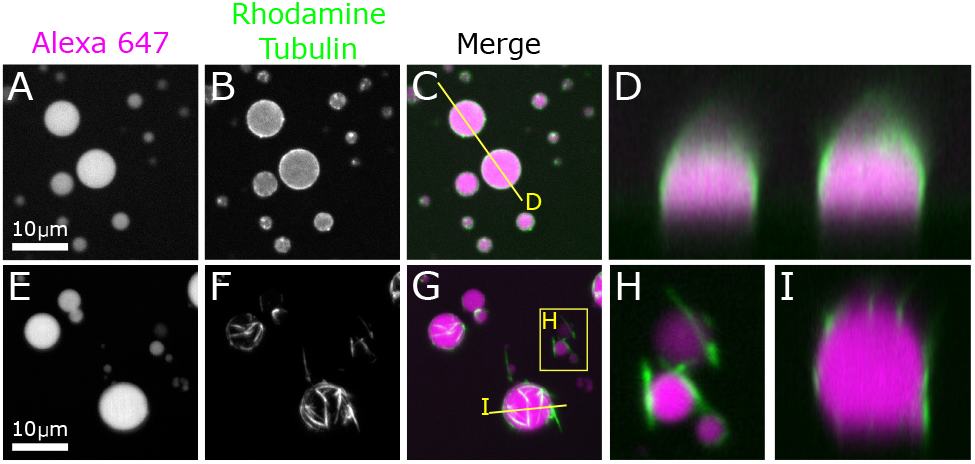
*β*-tubulin and microtubules localize to the surface of FUS-RNA droplets in *vitro*. (*A,B,C*) FUS-RNA droplets with tubulin sub-units. The droplet is visualized with Alexa 647 and tubulin with rhodamine. The mid-plane of a z-stack is shown. (*D*) Side-view of two droplets at the location indicated by the line in (*C*). (*E,F,G*) FUS-RNA droplets with taxol-stabilized microtubules. The maximum projection along z is shown to visualize all filaments around the droplets. (*H*) Zoomed image of two small droplets at the location indicated by the box in (*G*). (*I*) Side-view of a droplet at the location indicated by the line in (*G*).

Previous experiments and theory on the interaction of phase-separation and macromolecular networks have focused on the expansion of cavities in elastic networks [12–15, 18, 30, 44]. These deformations are driven by the free-energy per unit volume liberated by condensation, *i.e.* condensation pressure. For a single phase-separating component,

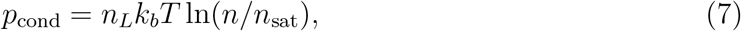

where *n_L_, n*, and *n*_sat_ are the number densities in the droplet, in the continuous phase, and at the point of saturation in the continuous phase, respectively [12]. When the surrounding medium is an ideal elastic solid, cavities grow without bound when *p_cond_* > 5*E*/6 [44]. While these ideas are well-suited for permanently cross-linked networks, visco-elastic relaxation of the cytoplasm [45–48] makes them largely irrelevant to the steady state of stress granules. Indeed, it is clear from Supplementary Movie 1 that stress granules and microtubules can re-arrange over timescales of a few minutes. Consequently, we expect elastic stresses induced by droplet growth to have fully relaxed within 90 minutes after induction of the granules, the focus of the current study.

Instead, the adhesion of tubulin and stress granules, as described above, appears to be the driving force underpinning the observed structural correlations of stress granules and tubulin. Generically, these surface affinities can do work when the *adhesion energy* is positive:

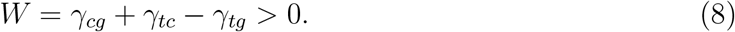

Since results for nocodazole-treated cells are consistent with |*γ_tc_* — *γ_tg_*| ≪ *γ_cg_*, we expect *W* ≈ *γ_cg_*. This positive adhesion energy could drive deformation of the microtubule network (Fig. 2) or stress granules (Fig. 3). Since the microtubule network presents little resistance to deformation on long time scales, we expect adhesive forces to consolidate the microtubule network around the droplet. This is consistent with macroscopic experiments and theory showing that neutrally wetting droplets preferentially sit at the side of filaments [49]. Because the persistence length of microtubules is large, concentration enhancements around stress granules must decay slowly (Fig. 2). Note that microtubules are too rigid to be significantly bent by sub-micron stress granules (see [50–53] and supplement section B12), while some bending is apparent in the adhesion of microtubules to larger *in vitro* droplets. Over long timescales, adhesion thus favors the migration of stress granules to regions of the cell with higher microtubule concentrations. In our experiments, we find that the microtubule-rich regions on either side of the nucleus are also the preferred location of stress granules (see Fig. 1 (*G,H*)). This effect is quantified by Fig. S8, where a voxel-wise cross-correlation of tubulin and G3BP1 intensities shows a strong tendency for granules to be found in regions of above average tubulin intensity.

We have introduced a statistical method to quantify structural correlations of various components in living cells, and identified significant interactions of microtubules and stress granules. This approach should be applicable to quantify a variety of weakly-bound structures within the cell. Depolymerization of microtubules and a physical model of adsorption support the hypothesis that non-specific adhesive interactions are sufficient to drive the observed structures. Tubulin sub-units’ weak affinity for the surface of stress granules, and weak repulsion from the bulk of stress granules, are amplified as microtubules polymerize, leading to the distinct enhancement of tubulin density around granules. With more precise structural data on the thickness of the interface between membraneless organelles as well as on the size of adsorbing particles, non-specific adsorption could provide a route to measuring *in vivo* surface tension through the fit parameter *a*_0_.

These interfacial phenomena could impact a wide-range of cellular phenomena. Surface partitioning could contribute to the observed core-shell structure of stress granules [9]. Wetting interactions of protein droplets with microtubules have been shown to facilitate branching [54, 55]. Because of their non-specific nature, we expect that interfacial forces could play a role in a number of interactions of membrane-less organelles with other supra-molecular structures. For example, the interaction of phase-separated domains with membrane strurctures including the endoplasmic reticulum [56], phagosome [57], Golgi apparatus [58], and synaptic vesicles [59]. We anticipate that similar interactions may also impact the localization of pathological liquid like aggregates as observed during neurodegenerative diseases [2, 60] or of functional aggregates in the formation of the skin barrier [61, 62].

## I. ACKNOWLEDGEMENTS

This work was primarily supported by grant numbers 172824 and 189940 from the Swiss National Science Foundation.

We thank Michael Murrell for helpful discussions and his micropatterning protocol. We thank Michael Steinmetz for advice and materials for *in vitro* experiments with tubulin. Moreover, we thank Mahdiye Ijavi for advice and discussion on *in vitro* experiments. We thank our lab members and members of the Bringing Materials to Life consortium at ETH for fruitful discussions.

## II. AUTHOR CONTRIBUTIONS

T.B, K.R., D.B., L.P. and E.D designed cell experiments; T.B and K.R. performed cell experiments; K.R., L.E., F.A., and E.D. designed *in vitro* experiments; K.R. performed *in vitro* experiments; Y.H. and L.E. purified protein for *in vitro* experiments; T.B. and E.D. developed statistical tools and analyzed data; T.B., R.S. and E.D. developed theoretical models; T.B. and E.D wrote the paper with input from all the authors.

## III. SUPPLEMENT

### A. Supplementary Movie

Epifluorescences time lapse of arsenite-treated U2OS cells. Tubulin in green and GSBP1 in magenta. Field of view is 26.3 × 16.4 *μ*m^2^, movies is played back at 210× the original speed

### B. Materials and Methods

#### 1. Cell culture

U2OS human osteosarcoma cells were grown in DMEM medium supplemented with 10% fetal bovine serum, and 2mM L-glutamine (all from Thermo Fisher), at 37°C in 5% CO_2_. For experiments, patterned coverslips were coated with 20 μg/ml of fibronection (Sigma) and single cells were plated on these coverslips 4—6 hours prior to experiments to ensure sufficient spreading.

#### 2. Micropatterning

Glass coverslips were incubated in 0.1 mg/ml poly-L-lysine-g-poly(ethyleneglycol) (PLL(20)-g[3.5]-PEG(2), SuSoS AG) for one hour. The coated coverslips were then exposed to deep UV light coated-side up in a UV/Ozone cleaner (ProCleaner Plus BioForce Nanosciences) through a chrome on quartz photomask (printed by Deltamasks). The transparent features on the otherwise opaque photomask are 25 μm-by-30 μm rectangles with two hemisphere caps of radius 12.5 μm at the ends and several hundred features per coverslip. Features are spaced at least 100 μm apart from edge to edge in all directions. After UV treatment, the patterned coverslips were coated with fibronectin and cells plated.

#### 3. Immunofluorescence

After specified durations of treatment with 0.5 mM sodium arsenite (Sigma) to induce stress granule formation (see e.g. [5]), wild-type U2OS cells were fixed with 4% formaldehyde for 15 min, permeabilized with 0.1% Triton X-100 in PBS, and blocked with 5 mg/ml bovine serum albumin (Sigma), in some cases containing also 0.25% Triton X-100. Fixed cells were incubated overnight at 4°C with primary antibodies: mouse anti-G3BP (1:500, abcam), rabbit anti-*β*-tubulin (1:200, abcam) to stain for stress granules and microtubules. Note that G3BP does not co-precipitate in *β*-tubulin immunoprecipitation and is commonly used as a stress granule marker [26, 63]. Secondary antibodies consisted of: Rhodamin Red-X anti-mouse IgG (1:500, Jackson ImmunoResearch) and Alexa 647 anti-rabbit IgG (1:500, Jackson ImmunoResearch). DNA has been stained for by incubating in DAPI (1:500, Sigma) for 15 min at room temperature. Coverslips were mounted in ProLong Gold (Thermo Fisher).

#### 4. Cell Imaging

Fixed cells were imaged on a Nikon Ti2 Eclipse with Yokogawa CSU-W1 spinning disk and 3i 3iL35 Laser Stack using a 100x oil objective with a numerical aperture of 1.45. The spatial resolution in the focal (xy-)plane was 0.065 μm/pixel and the step height (z-axis) was 0.2 μm. The theoretical diffraction limit for the shortest used wavelength (488 nm) is 130 nm, calculated as 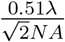 with numerical aperture *NA*, i.e. the full width half maximum of the theoretical point spread function of a confocal microscope [64]. For the actual optical setup, we assume a diffraction limit of three pixels (195 nm).

#### 5. Cell Detection and Sorting

Image analysis has been carried out using Matlab and has been automated to allow for efficient and reproducible processing of a large number of cells. Each set of confocal data acquired on the microscope contained one cell. The cell has been identified in the xy-plane using the REGIONPROPS function on an overlay of the maximum projections along *z* of all channels as well as a wide-field image of the nucleus through the DAPI stain. For the accurate detection of the cell shape in the focal plane, the image has been sharpen before thresholding. All images have been cropped around the detected cell in the *xy*-plane. To determine the *z*-coordinate of the base of each cell (cell base height) the sum of the median and 80th-percentile of each *xy*-plane for the filament channel has been collected. With only a small rim of background around the cropped cells, the median of the image is slightly below the median intensity of the cell and serves as a proxy for the overall structure of the cell. The 80th-percentile captures more pronounced features, such as individual filaments, while being robust against outliers. The cell base height has then been calculated as the *z*-coordinate corresponding to the maximal gradient of this intensity measure, which reliably detected the side of the cell attached to the coverslip. We assume the error of this measurement to be ±1 z-step. The maximum intensity z-slice was typically 2 to 4 slices above the cell base height. Cells in which this distance fell out of this interval were discarded. Moreover, cells with an area outside 85 to 110% of the area of the prescribed pattern (1257 μm^2^) were discarded. Cells with a ratio of the short principal axis to the long principal axis outside the interval [0.43 0.55] were also discarded. The corresponding ratio of the pattern itself is 0.45. These criteria have been chosen based on the corresponding histograms of all cells to discard outliers.

The background intensity of each channel has been approximated from the intensity in the corners of the cropped image outside the detected cell. Each channel has been corrected for this background intensity individually.

Multiple cells were recorded on each coverslip in one acquisition session. Within one such a batch, all patterns have the same orientation. The orientation of the patterns has been determined as the mean orientation of cells from one batch. Batches with less than 10 cells remaining after filtering the cell area and shape have been discarded. Individual cells with an orientation that deviates by more than 3° relative to the pattern orientation were also discarded. All cells have been rotated by the mean orientation of the pattern to ensure consistent cell orientation across samples. The remaining cells were aligned such that the centroid of the cell was in the center of the *xy*-plane. Because the nucleus is not always exactly in the center of the cell, cells were arranged such that the centroid of the nucleus always falls in the same hemisphere of the image, i.e. some cells were rotated by 180°. This yields cell stacks that are oriented in the xy-plane such that the long axis of the cell is horizontal and the nucleus falls to the left side.

#### 6. Intensity Normalization

The intensities of the G3BP1 and *β*-tubulin channel of each cell were individually normalized by their mean intensity 〈*I*〉 to account for differences in protein expression or staining. In order to define this mean intensity, we select a set of representative pixels. The x- and y-coordinates of representative pixels are those that fall inside the cell shape but outside the cell nucleus, based on the maximum projection along the z-coordinate of all channels. Pixel outside the cell or inside the nucleus were set to *NaN* (not a number) throughout all following analysis. The z-coordinates of the representative pixels are set as the second to fourth z-coordinate above the detected cell base height. Typically, the third z-coordinate above the cell base is the maximum intensity z-plane. The mean intensity is then calculated across all such representative pixels. Each channel is then normalized by the respective mean intensity in that cell.

#### 7. Stress Granule Detection

To analyse the interactions of stress granules with their surrounding, we need to first identify the granules. Stress granules were detected in the G3BP1 channel using an automated routine probing various global thresholds. This routine yields the spatial coordinates, centroid, volume and the eccentricity of well-defined stress granules within the bulk of the cytoplasm.

In a first step, the G3BP1 channel is filtered by subtracting a dynamic background. The background is determined by convolving a 30×30×5 box kernel on the confocal stack. The filtered image is then slightly blurred using the command *imgaussfilt* to decrease shot noise. The lowest probed threshold was set as the maximum of the mean intensity values in the xy-plane of the filtered G3BP1 channel across the height of the stack. The maximal threshold was set as the global maximum. 101 threshold values spaced evenly over the resulting range have been probed. For each threshold, the (matlab) function *regionprops3* has been used to detect volumetric blobs with intensity values above threshold. The number of detected blobs as well as median and mean volume have been recorded for each threshold. The aim is to detect stress granules, that is mathematically a group of finite size consisting of neighboring voxels. Further, the group of voxels must have a high intensity compared to the environment and be set apart by a reasonably sharp gradient in intensity to ensure that we do not detect fluctuations in the diffuse G3BP1 protein. Consequently, the optimal threshold shows a minimal change in the number of detected blobs upon varying the threshold level and corresponds to a local maximum in both median and mean volume, i.e. captures all or most granules in the cell. The mean volume ensures that we detect a large portion of the granules and do not overestimate the threshold. The median volume is sensitive to the distribution of sizes. Too low thresholds typically yield one large detected volume and many small blobs on the periphery. Thresholds that yield many small blobs will have a low median volume. The median volume consequently offers a handle to avoid underestimating the threshold. Weighting these three criteria (change in number of granules, mean and median volume) allows to assign a quality factor to each threshold and to detect a suitable value. Cells that do not exceed a minimal quality factor for the stress granule detection were discarded. In a second step, a logical three-dimensional mask of the detected granules, i.e. false where there is no granule and true for all voxels that are part of a granule, has been inflated by dilation with a sphere of radius 3 pixels. Applying these granule masks on the initial G3BP1 confocal stack, we can calculate the weighted centroid of the inflated and initially detected blobs. If the weighted centroids of a given blob differed by more than 0.5 pixels, the blob has also been discarded to ensure robustness of the detection. Moreover, all pairs of granules that fused upon dilation have been discarded to ensure a minimal distance between granules. Moreover, granules, typically very small, that reside below or above the cell nucleus were discarded as they fall outside the cell mask. The ellipticity of each granule has been defined as the ratio of major to minor principal axis of an ellipse fitted to each granule. Here, we only considered the principal axes in the xy-plane. The effective radius of the granule is defined as 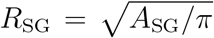, where *A*_SG_ is the area of the stress granule projected into the *xy*-plane like a shadow cast.

Aside from the characteristics (position, orientation, volume, etc.) a number of images are saved for each granule. All images are centered on the centroid of a given granule and show the xy-plane closest to the granule centroid. Note that pixels within these images that fall outside the cell limits or inside the respective cell nucleus are set to *NaN*. The size of these images is 221×221 pixels (about 14×14 μm), to capture also the long range deformations of the tubulin network around granules.

In the following analysis, granules with a radius below 3 pixels are discarded. Therefore, only granules with a diameter larger than twice the diffraction limit of 130nm are considered, see supplement section B4.

In a final step we filter granules after calculation of the individual distribution map in the G3BP1 channel (see e.g. Fig. 1 (*I*)). Granules that are not at least twice as bright as their local environment are discarded. The local environment is defined by dilating the individual granule shape with a disk of 8 pixel radius and subtracting the original granule shape. Further, granules with a mean relative intensity higher than the mean plus one standard deviation of the relative intensity across all granules are discarded to account for positive outliers.

#### 8. Construction of the reference cell

The alignment of all cells in three dimensions allows to construct cell stacks by overlaying all cells that belong to the same experimental conditions, e.g. duration of arsenite treatment. Each cell stack is blurred in the xy-plane using a Gaussian kernel with a variance of 4 pixels. While this method retains the overall intensity, noise as well as single filaments are blurred to suppress short range fluctuations. To capture the finite size of the cytosol as well as of the nucleus, each cell is masked by the cell outline and the nucleus shape. Based on these stacks, the reference intensity for any spatial coordinate is calculated as the mean intensity across all cells at this position, omitting the data from cells where the given pixel is masked. If more than half of the intensity values for a given coordinate fall within these masks of the nucleus or cell outline, the pixel is considered outside the cytoplasm of the reference cell. This way, the reference cell also serves as a mask for the expected cell shape (see Fig. 1 (*G*) and (*H*)).

Note that the symmetry along the short axis has deliberately been broken by rotating all cells such that the cell nucleus falls into the same half of the cell. The remaining symmetry of the pattern allows to fold the cell along the center line of the long axis to enhance the statistics for the calculation of the reference cell. Using *n* cells, this process yields 2*n* intensity values for all spatial coordinates in one hemisphere of the cell. The full reference cell is then recovered by mirroring the result along the center line.

Note that, due to fluctuations in the height of the cells, we only considered the intensity values between the cell base height up to 1.2 μm (6 slices) into the cell as reliable. Stress granules outside this range are not considered in the following analysis.

#### 9. Calculation of averaged images, distribution maps, radial distribution function g(r) and surface-relative distribution function g_s_(d)

To calculate averaged images we first define a subset of granules with comparable size and ellipticity. Images of round granules (i.e. with a principal axis ratio not exceeding 1.5), for example, are binned by the granule radius *R_SG_* in steps of three pixel (195 nm). Given a minimal granule radius of 195 nm, the first size bin is [195 nm 325 nm]. This size bin is then labeled as *R_SG_* = 0.26 μm. Unless stated otherwise, data are shown in bins that are statistically independent, i.e. each data point is assigned one bin.

To calculate the averaged images of a given bin (see e.g. Fig. 1 (*E,F*)), we average the intensity values for a given pixel location from all images of the G3BP1 and *β*-tubulin channel for granules of that bin. Note that we omit those pixels that fall outside the cytosol of the corresponding cell.

We calculate distribution maps (e.g. Fig. 1 (*K,L*) in the same way as averaged images only that each individual image is now normalized by the corresponding image of the reference cell. Normalization is done by point-wise division with the respective image originating from the reference cell of that channel at the same location. Note that, if a pixel is masked in either reference or G3BP1 and *β*-tubulin image, it is considered masked. Consequently, different pixels of a distribution map may not have the same number of pixels that contributed to the calculation of the corresponding intensity value.

Fluctuations in the number of contributing data sets for different pixel locations introduces a non-linearity when further analyzing distribution maps, which has to be taken into account when calculating the radial distribution *g*(*r*) of a distribution map. To capture the varying statistical weight, we do not take the radial average of the distribution map in question. Rather we take the average over all intensity values from the individually reference-cell-normalized images that the distribution map is calculated from at distance r from the center, again omitting masked entries. This way we know how many data points contribute to each entry in *g*(*r*). The error of *g*(*r*) at a given distance *r* is then calculated as the standard error, i.e. the standard deviation of the contributing values divided by the square root of the number of contributing values.

The surface-relative distribution function *g_s_*(*d*) is calculated from the reference-cell-normalized images for each granule individually, as it is the basis of the calculation of the partitioning coefficients for each granule. Here, the distance *d* for a given pixel location is calculated as the minimal distance to the outline of the stress granule. The outline itself has a value of *d* = 0 and pixels inside the granule have negative entries. The outline is defined by the stress granule detection routine. *g_s_*(*d*) is then calculated by averaging the intensities of pixels with the same *d*, omitting masked pixels. The error is again calculated as the standard error.

#### 10. Calculation of the partitioning coefficients k_g_ and k_s_

To calculate the partitioning coefficients *k_g_* = *c_g_/c_c_* and *k_s_* = *c_s_/c_c_*, we need to calculate the intensities of *β*-tubulin at the surface and in either bulk phase. Note that we assume that intensity and concentration are linearly proportional, the proportionality constant then cancels out when taking the ratio of intensities. Given that tubulin shows little preference to either bulk phase and is enhanced by a factor of less than 1.2 at the interface compared to the cytosol, we assume the same proportionality constant in and around a granule. In order to accurately access the intensity at the interface and around it, we base our calculation on the surface-relative distribution function *g_s_*(*d*). This way, variations in granule shape do not affect the measurement of surface intensity (see Fig. 5 (*A-C*)).

Based on *g_s_*(*d*) the intensity at the surface of a given granule *I_s_* is calculated as the maximum intensity in *g_s_*(*d*) within −2 to +2 pixels (−0.13 to 0.13 μm) around the maximal gradient ▽*g_s_*(*d*) of G3BP1, which corresponds to the position of maximum adsorption in the limit of neutral wetting of tubulin particles to the surface. The interval is chosen to allow for variations in the peak around the interface (see Fig. 4 (*C*)). At the surface, we find a broad peak of the tubulin intensity, with a width typically larger than the point-spread-function of the microscope (195 nm), suggesting that, while optical artifacts are certainly expected to affect the shape of the peak, the peak is indeed broad. Note that the width of the peak coincides with the region over which the intensity of G3BP1 transitions from bulk cytosol to bulk granule. In order to avoid artifacts from this broad peak across the interface of the stress granule, we measure the intensity inside the granule *I_g_* as the mean of *g_s_*(*d*) inside the granule for *d* ≤ 0.39 μm from the detected surface. We also omit the entry in *g_s_*(*d*) with the largest distance from the surface, which corresponds to the very center and typically has poor statistics. To have a reasonable set of pixels to average over, we only consider granules with a radius larger than 0.59 μm for this analysis, i.e. at least three entries with *d* ≤ 0.39 μm.

We have previously shown that granules, also in nocodazole treated cells, typically reside in a region of higher than average tubulin intensity (see Fig. 1 (*G,H*) and Supplementary Fig. S7). We can therefore not just take the expected tubulin intensity of the reference cell (which is one by construction), but also have to define an intensity for the surrounding of each granule *I_c_*. In nocodazole-treated cells, the long range decay of tubulin around granules is significantly less pronounced compared to cells with intact microtubule networks (see Fig. 4), but still present. The intensity of the surrounding is thus depending on the distance from the granule and consequently subject to error. To capture the tubulin density the granule experiences, we measure as close to the granule as possible while avoiding artifacts from the broad surface peak in tubulin localized to the interface zone of the granule. We chose to define the intensity in the surrounding as the mean of the *g_s_*(*d*) between d = 0.46 and *d* = 0.59 μm outside the granule, keeping the same 0.39 μm away from the surface as for the calculation of *I_g_*. This half width of the interface of 0.39 μm is chosen to be broad enough to also accommodate error from the position of the maximum gradient, which, on average, is found at *d* = —1.4 ± 1 pixels and thus does not coincide with *d* = 0.

#### 11. Fitting

Fit parameters and errors are evaluated through variation of the fit over *N* = 1000 realizations. From the initial fit of the theory (Eq. 5) to the data, we extract the residuals and initial fit parameter *a*_0_. Using this fit parameter, we assign each data point with the theoretical prediction of —Δ*G_s_*(—Δ*G_g_*), based on the — Δ*G_g_* of the data. For each iteration, we then add Gaussian noise to the data with zero mean and variance equal to the standard deviation of the residuals to create a variance of the original data and perform a new fit. The reported value of *a*_0_ is the mean of *N* = 1000 realizations of the variation, with the error given as the standard deviation. All fits use bisquare weighting of residuals.

#### 12. Calculation of the bendocapillary length

To estimate the stress granule radius at which we can expect microtubules to bend around the granule we perform a gedankenexperiment where a microtubule wraps a stress granule once. The elastic energy to bend a length *l* of filament to a radius of curvature matching the radius of the stress granule *R_SG_* is then

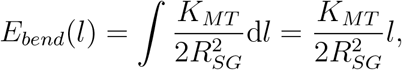

With bending rigidity of the microtubule *K_MT_* and line integral *∫ dl*. We can approximate the *K_MT_* from the persistence length *l_p_* of microtubules, which we take to be one millimeter:

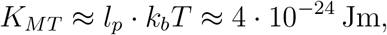

with thermal energy *k_b_T*. The adhesion energy of a microtubule of length *l* in contact with the granule is then

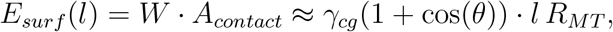

with surface tension *γ_cg_* between granule and cytosol and contact angle *θ* between granule and microtubule and microtubule radius *R_MT_*. Typical values of the surface tension of protein droplets vary between 1 to 100 μJ/m^2^ [38, 65]. For the stress granules, classical wetting, that is assuming a thin interface of the granule, suggests a maximum value of the surface tension of about 10^-5^ J/m^2^. Equating both energies we get a granule radius *R_bc_* at which both forces are equal in the range of

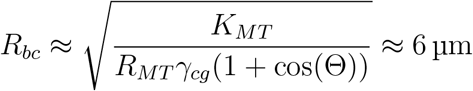

assuming neutral wetting, a surface tension of 10 5 J/m^2^ and a microtubule radius of *R_MT_* ≈ 12 nm. Note that lower surface tension leads to larger *R_bc_*.

#### 13. Droplets in vitro

His- and gb1-tagged full length FUS protein was purified from *E. coli* under denaturing conditions with affinity chromatography. Overnight incubation with TEV protease and subsequent second affinity column removed the tags. Protein was concentrated in presence of 6M urea up to 1.5mM, similar to FUS NTD protocol [66]. Full length FUS was purified from *E. coli* and concentrated up to 1.5mM in the presence of 6M urea. To induce phase separation, the stock protein solution was diluted to a final concentration of 50 μM in buffer (50mM HEPES, 150mM NaCl, pH 7.5). RNA was added to the FUS solution in the form of poly-U (Sigma Aldrich) at a final concentration of 0.5mg/ml. To image the droplets in fluorescence, 1 *μ*M AlexaFluor 647 (ThermoFisher Scientific) was mixed into the droplet solution, where it partitioned into the condensed phase.

10% rhodamine-tagged porcine tubulin (Cytoskeleton, Inc.) was mixed with purified bovine tubulin in tubulin buffer (80mM PIPES, 1mM EGTA, 2mM MgCl_2_, pH 6.9). This was either added to the droplets as sub-units at a final concentration of 0.1 mg/ml, or first polymerized in the presence of 1mM GTP and stabilized with Taxol (Paclitaxel, ThermoFisher Scientific). A small amount of the resulting microtubules were then added to the droplet solution.

Directly after mixing, droplets were pipetted between sandwiched coverslips containing a coating of oil with 2% FluoroSurfactant (RAN Biotechnologies). The coverslips were sealed and imaged with a 60x water immersion lens (NA = 1.2).

### C. Theory: Partition Coefficients

To evaluate under which conditions a tubulin sub-unit will prefer to reside on the granule surface or in bulk cytosol or bulk granule, we identify the chemical potential *μ* for each state assuming tubulin is dilute:

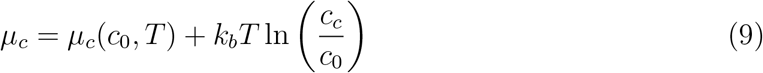

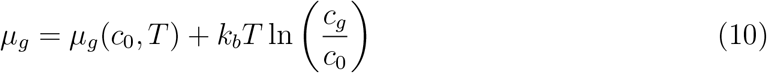

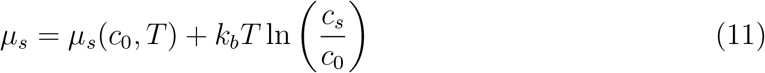

Note that the surface concentration *c_s_* is also a volumetric concentration. This is because the tubulin sub-units have a volumetric extend, and we measure intensity per voxel in the experiment. Moreover, the surface is not as well defined as, for example, an air-water interface under room temperature and normal pressure. This leads to a surface phase with a certain extend.

We assume a local equilibrium of the chemical potential of tubulin in and around the granule. We justify this for once, because we consider a small subset of the cell, i.e. inside and outside of the granule are close together, and because we assume that tubulin sub-units are inert with respect to the granule. Equating the chemical potentials of a sub-unit in the cytosol and inside the granule, we can define the partition coefficient between granule and cytosol

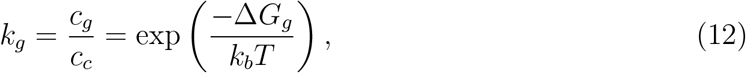

with free energy difference — Δ*G_g_* = *μ_c_*(*c*_0_,*T*) — *μ_g_*(*c*_0_,*T*) between a sub-unit in the granule compared to the cytosol. Analogously, we define the partitioning coefficient to the surface relative to the cytosol as

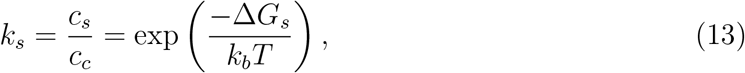

with free energy difference — Δ*G_s_* = *μ_c_*(*c*_0_,*T*) — *μ_s_*(*c*_0_,*T*).

In order to calculate the free energy differences Δ*G_g_* and Δ*G_s_*, we present two models, which differ in the definition of the interface between stress granule and cytosol. The tubulin sub-unit is, in both cases modeled as a colloidal particle.

**FIG. S1.**
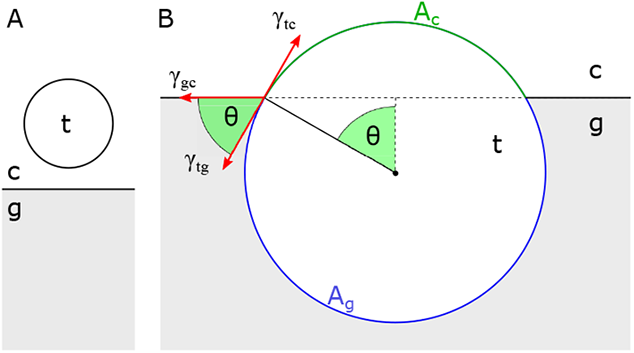
Tubulin sub-unit modelled as a colloidal particle *t* interacting with a thin interface between stress granule *g* and cytosol *c*. (*A*) Reference state of a sub-unit with surface area *A*_0_ in bulk cytosol. (*B*) A sub-unit wetting the interface with contact angle *θ*. *A_c_* is the surface area of the spherical cap of the sub-unit exposed to the cytosol, *A_g_* = *A*_0_ — *A_c_* respectively the spherical cap exposed to the granule.

### D. Theory: Thin interface

First, let us consider the colloidal model tubulin particle of radius *R* interacting with an interface of width 2*w* that is much thinner than the size of the particle *R* ≫ *w*. (Note that the smallest possible tubulin particle is a sub-unit but we cannot ensure that all tubulin particles are sub-units in nocodazole-treated cells.) This corresponds to the classical picture of a particle at an interface as shown in Fig. 5. — Δ*G_g_* = *μ_c_*(*c*_0_,*T*) — *μ_g_*(*c*_0_,*T*) is the energy difference a sub-unit experiences when it is moved from bulk cytosol to inside the granule. Note that the assumption that tubulin is dilute implies that tubulin sub-units do not feel and interact with each other. Δ*G_g_* is then given as the surface area of the sub-unit *A*_0_ times the difference in the surface tension of the sub-unit towards the cytosol *γ_tc_* and towards the bulk of the granule *γ_tg_*

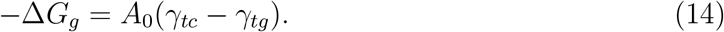

In order to calculate the free energy difference — Δ*G_s_* = *μ_c_*(*c*_0_,*T*) — *μ_s_*(*c*_0_,*T*) between a sub-unit in bulk cytosol and adhered to the surface of the granule, let us first consider the energy of a sub-unit at the interface. This energy is given as the balance of the area of the interface taken up by the sub-unit *A_cut_* times the surface tension of the granule towards the cytosol *γ_cg_* and the surface areas *A_c_* and *A*_0_ — *A_c_* of the granule exposed to the cytosol and the granule multiplied with the respective surface tension:

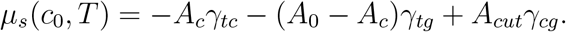

The energy difference between a sub-unit in bulk cytosol and on the surface is then

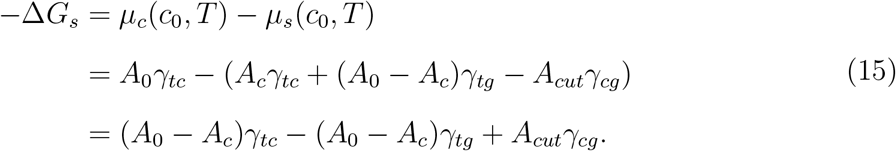

Using Young’s law

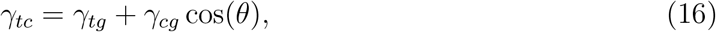

with contact angle *θ*, we find

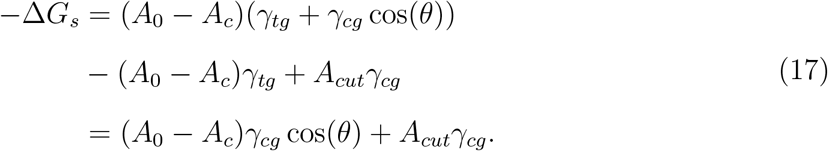

To define the geometric quantities, we now consider a spherical sub-unit, as illustraded in Fig. S1. The sub-unit with radius *R* then has a surface area *A*_0_ = 4*πR*^2^, surface area exposed to the cytosol (in the shape of a spherical cap) of *A_c_* = 2*πR*^2^(1 — cos(*θ*)) and an area *A_cut_* = *πR*^2^ sin^2^(*θ*) that the sub-unit takes up on the surface of the granule, with contact angle *θ* as defined in Fig. 5 (*B*). Thus we finally arrive at

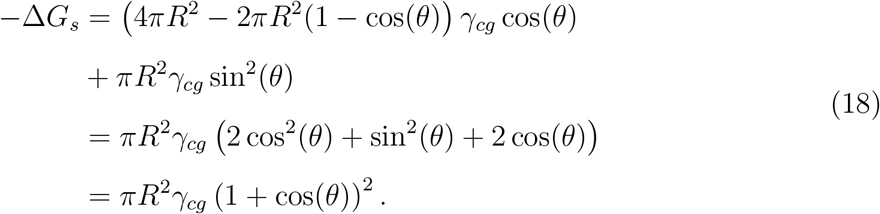

The energy difference between a particle in bulk cytosol compared to bulk granule is then analogously

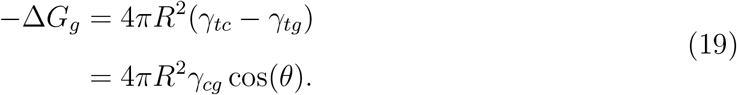

For comparison to our data, we express Δ*G_s_* in terms of Δ*G_g_*

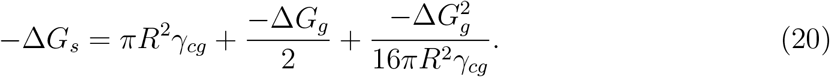

Note that this expression has removed the explicit dependence on contact angle. Δ*G_g_* is a function of both *γ_cg_* and *θ*. The histogram of observed Δ*G_g_* (Fig. 5 (*D*)), however, shows both positive and negative values with a mean close to zero. If *θ* was constant, this would call for negative surface tension *γ_cg_*, which is not physical. We thus assume that variations in Δ*G_g_* are dominated by variations in contact angle *θ* and hold *γ_cg_* constant.

### E. Theory: Thick interface

Tubulin sub-units, as well as the constituents of stress granules, are proteins, i.e. the assumption that the particle interacting with a droplet interface is much larger than the interface width is not given. Moreover, our data suggest a width of the interface of stress granules of *w* ≈ 0.4 μm, much larger than a tubulin sub-unit with radius *R* ≈ 3 nm [37]. To set up a theory for a thick interface (*R* ≪ *w*), we follow the approach presented by Cahn and Hillard [39]. We consider a two-phase system characterized by an intensive scalar quantity *ϕ* (other than temperature or pressure) that transitions smoothly from one state (inside the bulk of the stress granule) to another state (bulk cytosol). Here we express *ϕ* as the local mesoscale composition

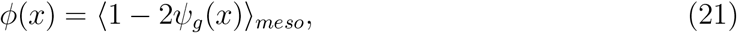

where *ψ_g_* gives the local volume fraction of stress granule components and 〈·〉_*meso*_ gives the local average over a volume sufficiently large such that *ϕ* is smooth [41]. The spatial coordiante *x* is normal to a flat interface between granule and cytosol, with the midpoint of the interface at *x* = 0, without loss of generality.

We assume a Helmholtz free energy per unit volume of the system given as

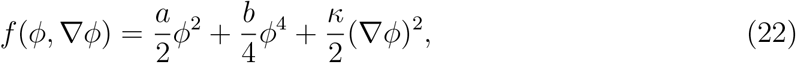

with parameter *a* = *a*(*T*) that is negative under conditions in which the system separates and positive constants *b* and *κ* [39-41]. The volume terms capture the phase behavior of the system and the gradient term describes interfacial energies. For negative *a*, i.e. below the critical temperature, *f* has two minima at 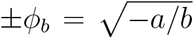 [41]. Assuming a constant chemical potential *μ* everywhere in the system, it can be shown that the local composition takes the form [40, 41]

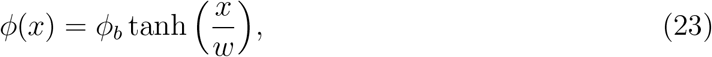

with width of the interface [40]

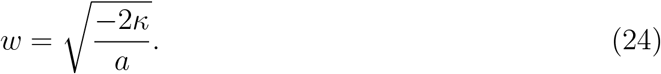

**FIG. S2.**
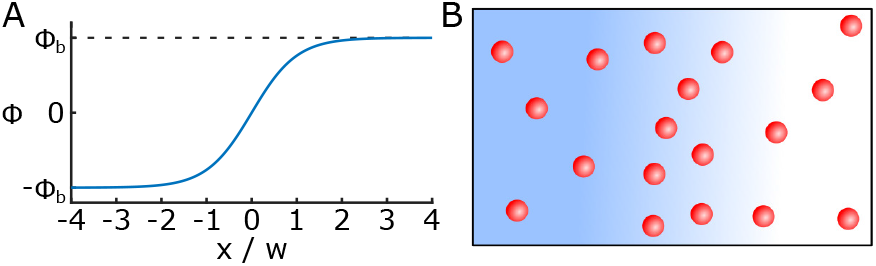
Tubulin sub-units interacting with an extended interface between granule and cytosol. (*A*) Phases and interface are determined by 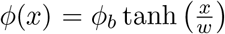. (*B*) Tubulin sub-units (red) are smaller than the interface width 2*w*.

Integrating the free energy over the interface, one finds the surface tension between both phases [41]

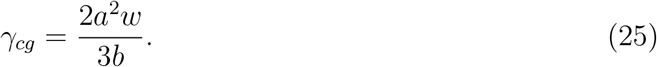

To evaluate the energy per unit volume around the interface, we expand *ϕ*(*x*) around *x* = 0

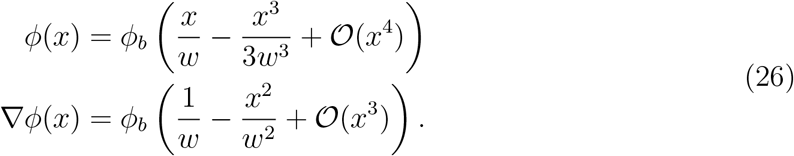

Inserting into equation 22 we find

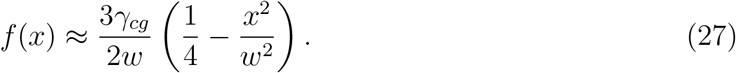

Far from the interface, we find for either bulk phase

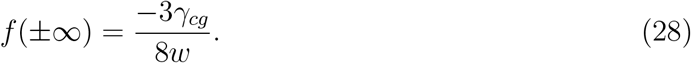

Moving a particle with surface area *A*_0_ and volume *V*_0_ from bulk cytosol (*x* → ∞) to position *x* we then change the free energy by

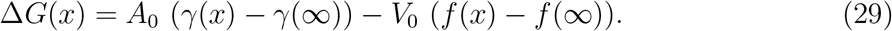

Δ*G*(*x*) consists of a contribution from the surface tension of the particle proportional to its surface area and a term that captures the energy of the displaced fluid. For the change in surface tension we apply the law of mixtures and find

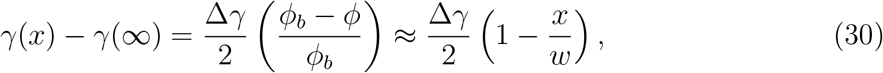

with Δ*γ* = *γ_tg_* — *γ_tc_*, where the last term shows the expansion around *x* = 0. For the free energy difference between the bulk phases, the volume terms cancel out (see eq. 28) and we find

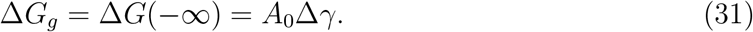

Inserting equations 27, 28 and 30 into equation 29, we can express the energy of a particle around the interface as

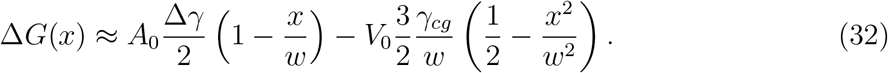

Δ*G*(*x*) has a minimum at the equilibrium position for the particle, for which we find

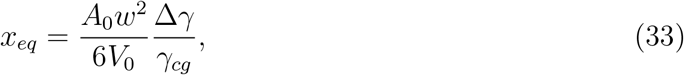

with energy

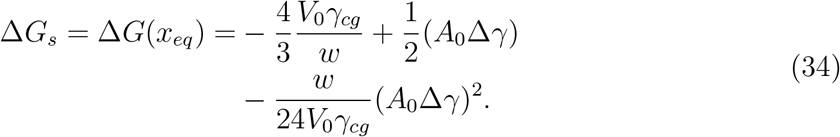

Here we can identify Δ*G_g_* and arrive at

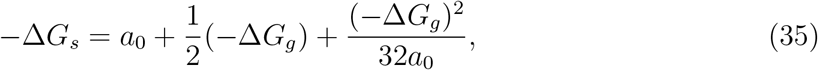

with parameter

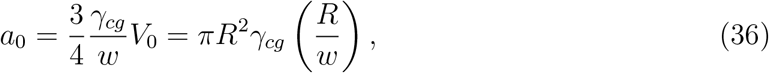

assuming a spherical particle with radius *R*. Note that the theory predicts a broad adsorption peak with a width corresponding to the width of the interface.

**FIG. S3.**
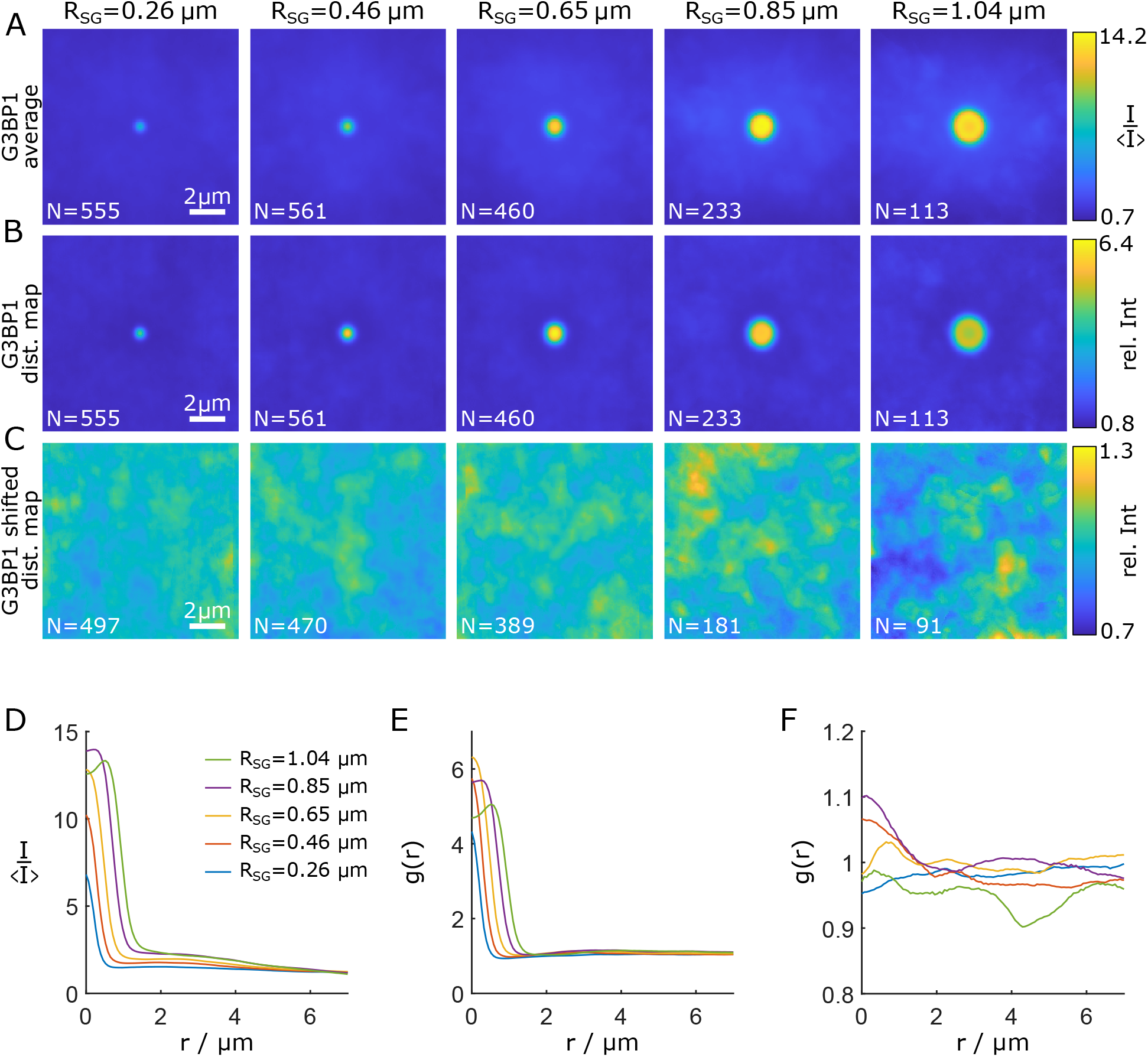
Average images, and distribution maps of stress granules. (*A*) Average images of G3BP1 for different granule sizes, *N* gives the number of contributing granules. (*B*) Distribution maps of G3BP1. (*C*) Negative control: Distribution maps of G3BP1 where images of granule in cell *i* have been extracted at the same location in cell *i* + 1, probing random but biologically plausible input. Note that granules with locations that fall within the cell nucleus or outside the cell at cell *i* + 1 are discarded. (*D*) Radial curves corresponding to the average images in (*A*). (*E*) *g*(*r*) corresponding to the distribution maps in (*B*). (*F*) *g*(*r*) corresponding to the distribution maps in (*C*).

**FIG. S4.**
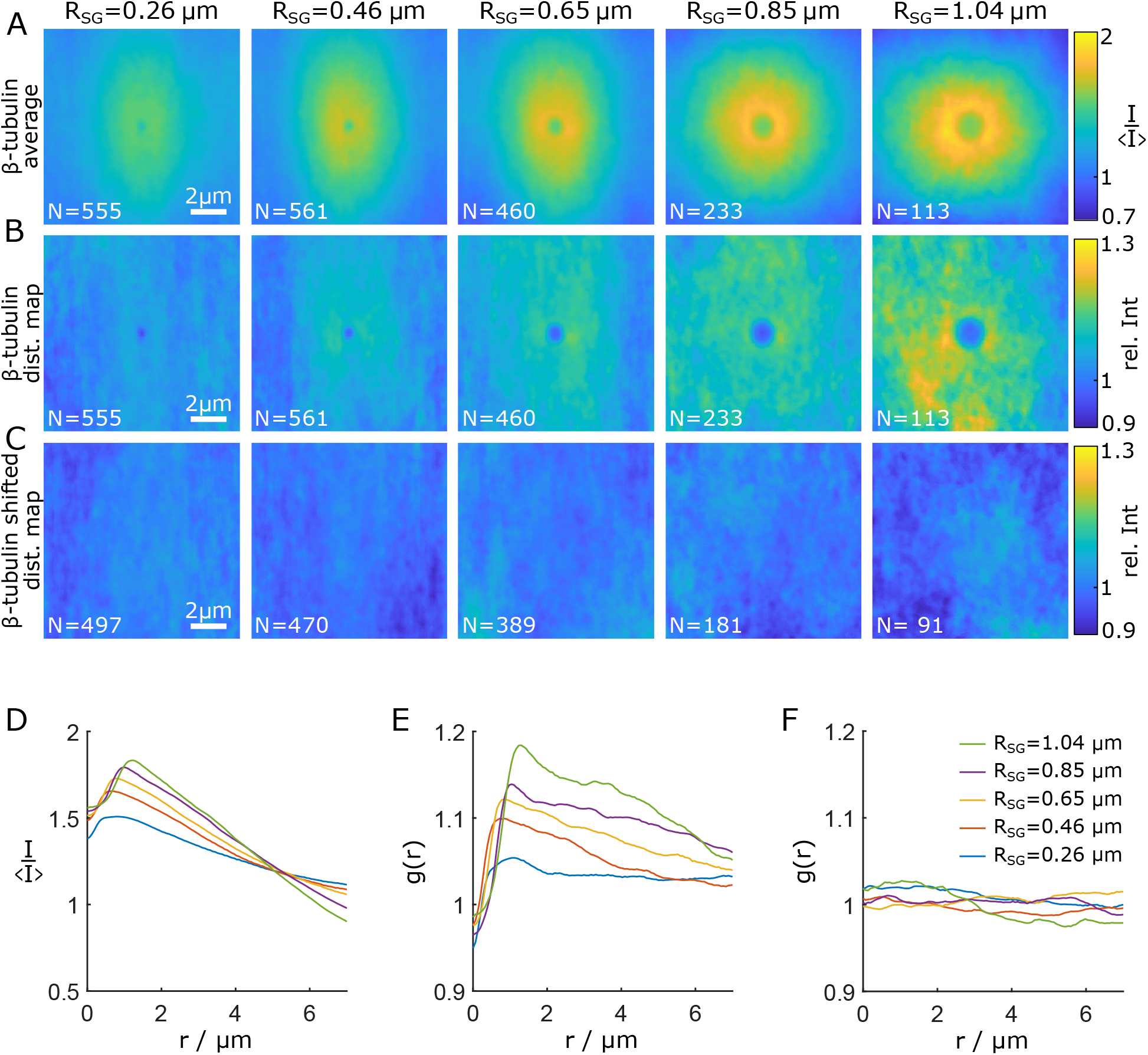
Average images, and distribution maps of *β*-tubulin. (*A*) Average images of *β*-tubulin for different granule sizes, *N* gives the number of contributing granules. (*B*) Distribution maps of *β*-tubulin. (*C*) Negative control: Distribution maps of *β*-tubulin where images of the *β*-tubulin channel in cell *i* have been extracted at the same location in cell *i* + 1, probing random but biologically plausible input. Note that images with locations that fall within the cell nucleus or outside the cell at cell *i* + 1 are discarded. (*D*) Radial curves corresponding to the average images in (*A*). (*E*) *g*(*r*) corresponding to the distribution maps in (*B*). (*F*) *g*(*r*) corresponding to the distribution maps in (*C*).

**FIG. S5.**
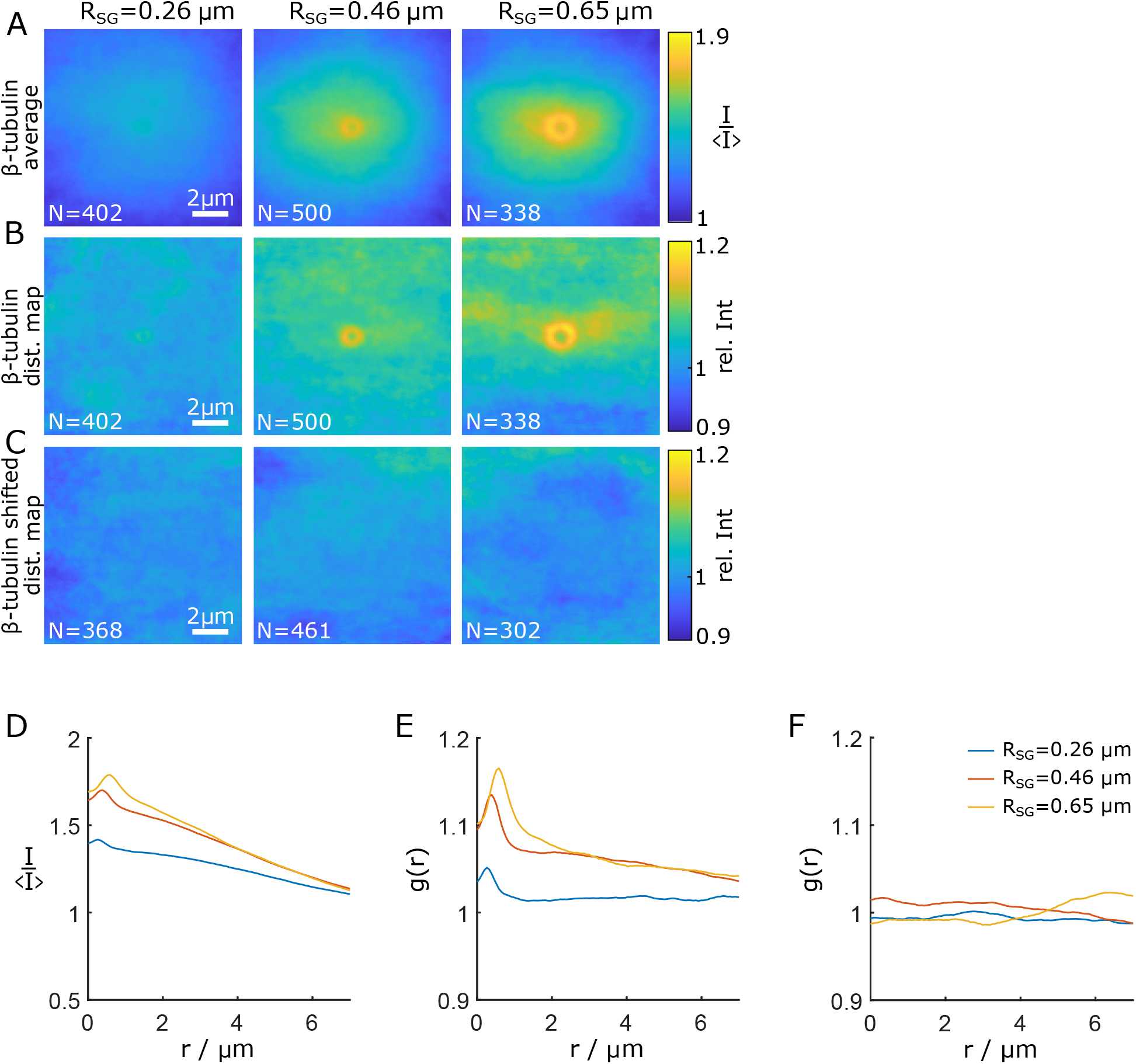
Average images, and distribution maps of *β*-tubulin in nocodazole-treated cells. (*A*) Average images of *β*-tubulin for different granule sizes, *N* gives the number of contributing granules originating from nocodazole-treated cells. (*B*) Distribution maps of *β*-tubulin in nocodazole-treated cells. (*C*) Negative control: Distribution maps of *β*-tubulin where images of the *β*-tubulin channel in cell *i* have been extracted at the same location in cell *i* + 1, probing random but biologically plausible input from nocodazole-treated cells. Note that images with locations that fall within the cell nucleus or outside the cell at cell *i* + 1 are discarded. (*D*) Radial curves corresponding to the average images in (*A*). (*E*) *g*(*r*) corresponding to the distribution maps in (*B*). (*F*) *g*(*r*) corresponding to the distribution maps in (*C*).

**FIG. S6.**
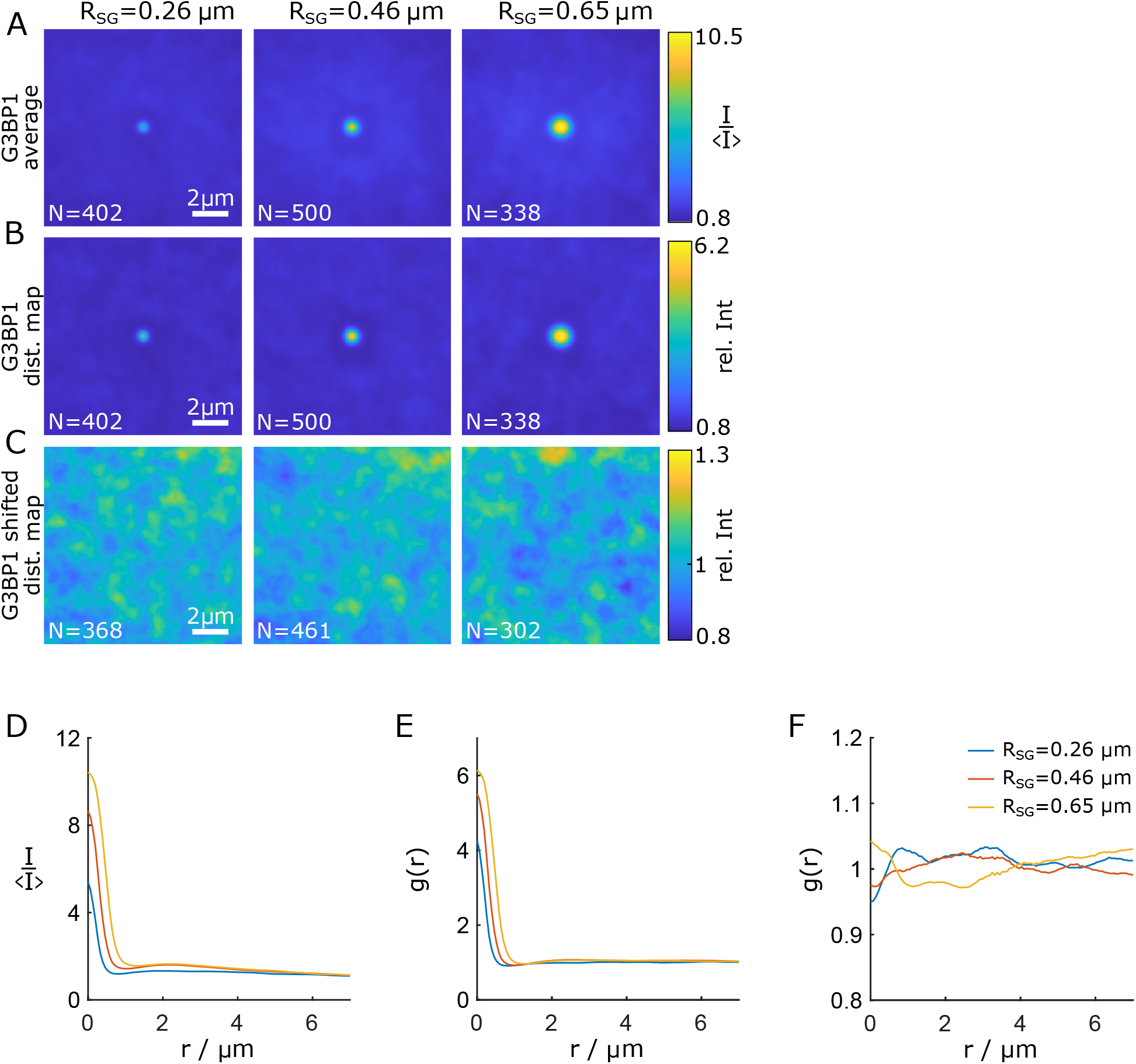
Average images, and distribution maps of G3BP1 in nocodazole-treated cells. (*A*) Average images of G3BP1 for different granule sizes, *N* gives the number of contributing granules originating from nocodazole-treated cells. (*B*) Distribution maps of G3BP1 in nocodazole-treated cells. (*C*) Negative control: Distribution maps of G3BP1 where images of the *β*-tubulin channel in cell *i* have been extracted at the same location in cell *i* + 1, probing random but biologically plausible input from nocodazole-treated cells. Note that images with locations that fall within the cell nucleus or outside the cell at cell *i* + 1 are discarded. (*D*) Radial curves corresponding to the average images in (*A*). (*E*) *g*(*r*) corresponding to the distribution maps in (*B*). (*F*) *g*(*r*) corresponding to the distribution maps in (*C*).

**FIG. S7.**
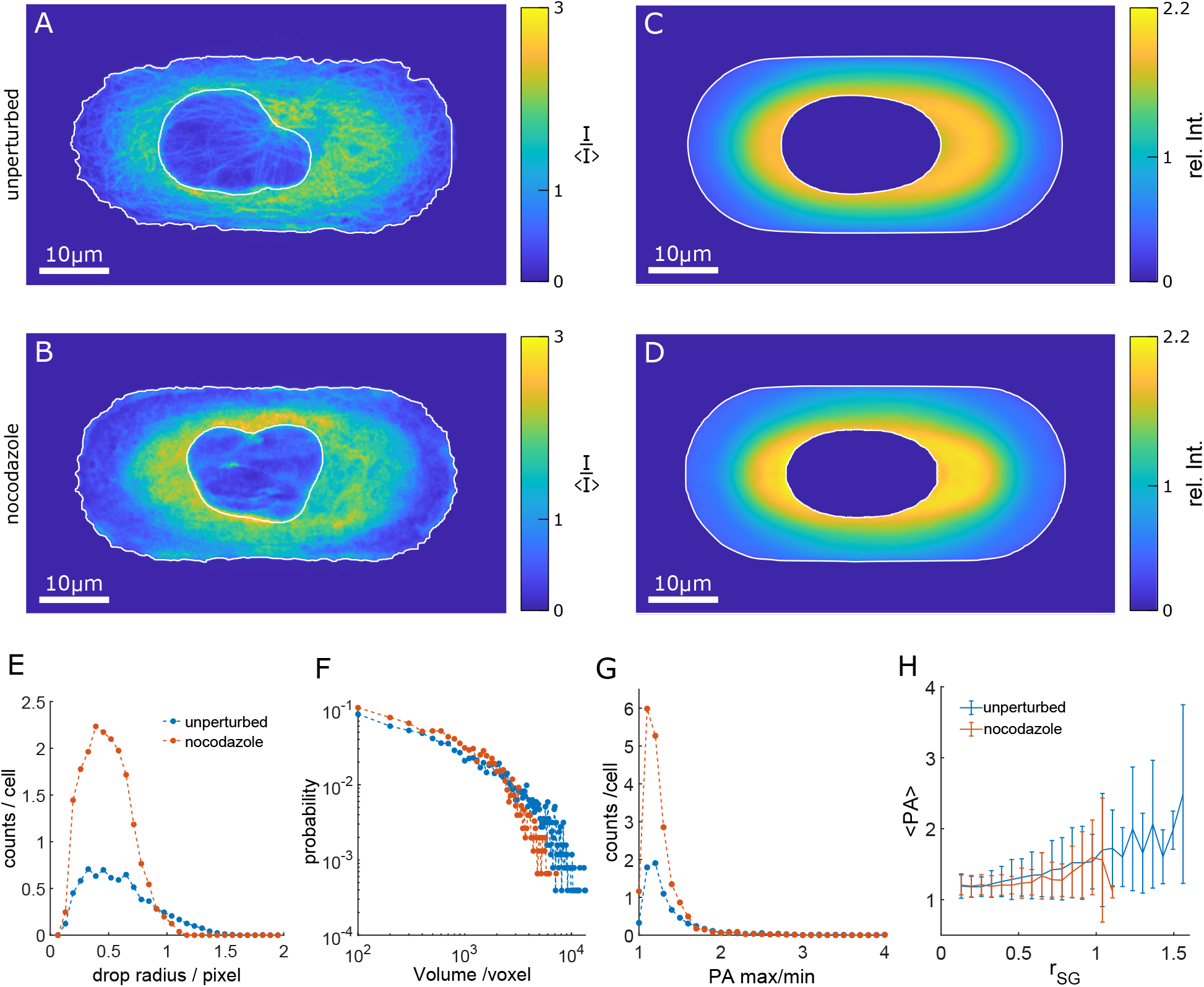
Comparison of cells with and without nocodazole treatment (*A*) xy-slice of the tubulin channel of an exemplary cell with intact microtubule network. Cell and nucleus outline are shown in white. (*B*) xy-slice of the tubulin channel of an exemplary cell after 90 minutes of nocodazole treatmeant. Cell and nucleus outline are shown in white. (*C*) xy-slice of the tubulin reference cell with intact microtubule network. Average cell and nucleus outline are shown in white. (*D*) xy-slice of the tubulin reference cell after 90 minutes of nocodazole treatmeant. Average cell and nucleus outline are shown in white. (*E*) Histogram of observed granule radii with a total of 2543 granules from 335 cells with intact microtubule network and 1520 granules from 81 cells with nocodazole treatment. (*F*) Histogram of granule volume. (*G*) Histogram of the principal axis ratio. (*H*) Mean principal axis ratio as a function of granule radius. The errorbars are given as the standard deviation.

**FIG. S8.**
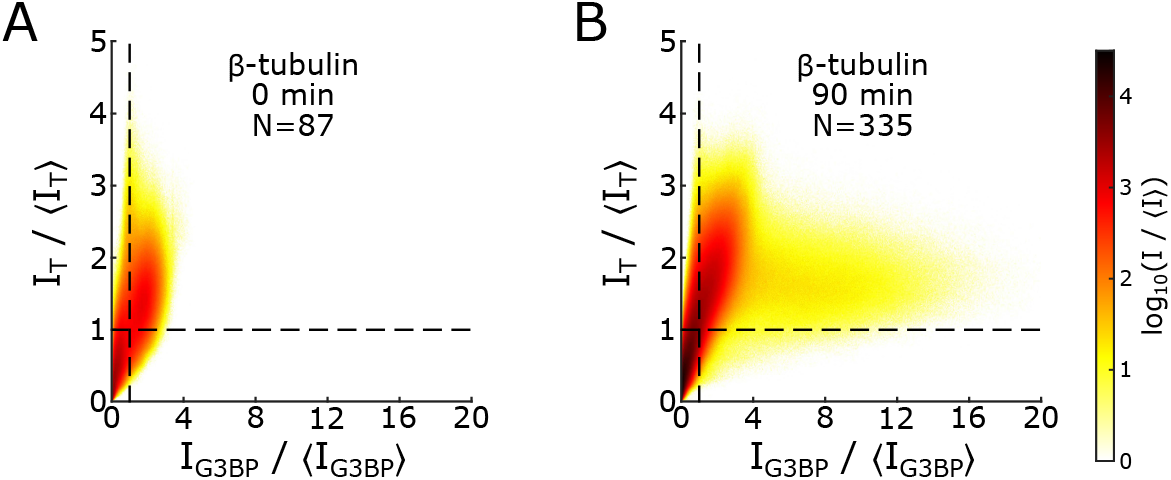
Spatial correlation between tubulin and G3BP1 in the reference cells. (*A*) For cells without arsenite treatment (0minutes of arsenite treatment). (*B*) After 90 minutes of arsenite treatment. 〈·〉 gives the average intensity of the respective channel.

## References

[1] S. F. Banani, H. O. Lee, A. A. Hyman, and M. K. Rosen, “Biomolecular condensates: Organizers of cellular biochemistry,” Nature Reviews Molecular Cell Biology, vol. 18, pp. 285–298, May 2017.

[2] Y. Shin and C. P. Brangwynne, “Liquid phase condensation in cell physiology and disease,” Science, vol. 357, p. eaaf4382, Sept. 2017.

[3] F.-M. Boisvert, S. van Koningsbruggen, J. Navascués, and A. I. Lamond, “The multifunctional nucleolus,” Nature Reviews Molecular Cell Biology, vol. 8, pp. 574–585, July 2007.

[4] J. R. Buchan and R. Parker, “Eukaryotic Stress Granules: The Ins and Outs of Translation,” Molecular Cell, vol. 36, pp. 932–941, Dec. 2009.

[5] N. Kedersha, G. Stoecklin, M. Ayodele, P. Yacono, J. Lykke-Andersen, M. J. Fritzler, D. Scheuner, R. J. Kaufman, D. E. Golan, and P. Anderson, “Stress granules and processing bodies are dynamically linked sites of mRNP remodeling,” The Journal of Cell Biology, vol. 169, pp. 871–884, June 2005.

[6] C. P. Brangwynne, C. R. Eckmann, D. S. Courson, A. Rybarska, C. Hoege, J. Gharakhani, F. Julicher, and A. A. Hyman, “Germline P Granules Are Liquid Droplets That Localize by Controlled Dissolution/Condensation,” Science, vol. 324, pp. 1729–1732, June 2009.

[7] C. P. Brangwynne, T. J. Mitchison, and A. A. Hyman, “Active liquid-like behavior of nucleoli determines their size and shape in Xenopus laevis oocytes,” Proceedings of the National Academy of Sciences, vol. 108, pp. 4334–4339, Mar. 2011.

[8] J. Berry, S. C. Weber, N. Vaidya, M. Haataja, and C. P. Brangwynne, “RNA transcription modulates phase transition-driven nuclear body assembly,” Proceedings of the National Academy of Sciences, vol. 112, pp. E5237–E5245, Sept. 2015.

[9] S. Jain, J. R. Wheeler, R. W. Walters, A. Agrawal, A. Barsic, and R. Parker, “ATPase-Modulated Stress Granules Contain a Diverse Proteome and Substructure,” Cell, vol. 164, pp. 487–498, Jan. 2016.

[10] V. Marx, “Cell biology befriends soft matter physics,” Nature Methods, vol. 17, pp. 567–570, June 2020.

[11] T. Wiegand and A. A. Hyman, “Drops and fibers — how biomolecular condensates and cytoskeletal filaments influence each other,” Emerging Topics in Life Sciences, vol. 4, pp. 247–261, Dec. 2020.

[12] R. W. Style, T. Sai, N. Fanelli, M. Ijavi, K. Smith-Mannschott, Q. Xu, L. A. Wilen, and E. R. Dufresne, “Liquid-Liquid Phase Separation in an Elastic Network,” Physical Review X, vol. 8, p. 011028, Feb. 2018.

[13] K. A. Rosowski, T. Sai, E. Vidal-Henriquez, D. Zwicker, R. W. Style, and E. R. Dufresne, “Elastic ripening and inhibition of liquid-liquid phase separation,” Nature Physics, Jan. 2020.

[14] J. Y. Kim, Z. Liu, B. M. Weon, T. Cohen, C.-Y. Hui, E. R. Dufresne, and R. W. Style, “Extreme cavity expansion in soft solids: Damage without fracture,” Science Advances, vol. 6, p. eaaz0418, Mar. 2020.

[15] Y. Shin, Y.-C. Chang, D. S. Lee, J. Berry, D. W. Sanders, P. Ronceray, N. S. Wingreen, M. Haataja, and C. P. Brangwynne, “Liquid Nuclear Condensates Mechanically Sense and Restructure the Genome,” Cell, vol. 175, pp. 1481–1491.e13, Nov. 2018.

[16] H. D. Ou, S. Phan, T. J. Deerinck, A. Thor, M. H. Ellisman, and C. C. O’Shea, “ChromEMT: Visualizing 3D chromatin structure and compaction in interphase and mitotic cells,” Science, vol. 357, p. eaag0025, July 2017.

[17] C. M. Caragine, S. C. Haley, and A. Zidovska, “Nucleolar dynamics and interactions with nucleoplasm in living cells,” eLife, vol. 8, p. e47533, Nov. 2019.

[18] D. S. W. Lee, N. S. Wingreen, and C. P. Brangwynne, “Chromatin mechanics dictates subdiffusion and coarsening dynamics of embedded condensates,” Nature Physics, Jan. 2021.

[19] A. Aizer, Y. Brody, L. W. Ler, N. Sonenberg, R. H. Singer, and Y. Shav-Tal, “The Dynamics of Mammalian P Body Transport, Assembly, and Disassembly In Vivo,” Molecular Biology of the Cell, vol. 19, pp. 4154–4166, Oct. 2008.

[20] D. W. Sanders, N. Kedersha, D. S. Lee, A. R. Strom, V. Drake, J. A. Riback, D. Bracha, J. M. Eeftens, A. Iwanicki, A. Wang, M.-T. Wei, G. Whitney, S. M. Lyons, P. Anderson, W. M. Jacobs, P. Ivanov, and C. P. Brangwynne, “Competing Protein-RNA Interaction Networks Control Multiphase Intracellular Organization,” Cell, vol. 181, pp. 306–324.e28, Apr. 2020.

[21] N. L. Kedersha, M. Gupta, W. Li, I. Miller, and P. Anderson, “RNA-Binding Proteins Tia-1 and Tiar Link the Phosphorylation of Eif-2α to the Assembly of Mammalian Stress Granules,” The Journal of Cell Biology, vol. 147, pp. 1431–1442, Dec. 1999.

[22] A. Aulas, M. M. Fay, S. M. Lyons, C. A. Achorn, N. Kedersha, P. Anderson, and P. Ivanov, “Stress-specific differences in assembly and composition of stress granules and related foci,” Journal of Cell Science, vol. 130, pp. 927–937, Mar. 2017.

[23] E. Kolobova, A. Efimov, I. Kaverina, A. K. Rishi, J. W. Schrader, A.-J. Ham, M. C. Larocca, and J. R. Goldenring, “Microtubule-dependent association of AKAP350A and CCAR1 with RNA stress granules,” Experimental Cell Research, vol. 315, pp. 542–555, Feb. 2009.

[24] K. G. Chernov, A. Barbet, L. Hamon, L. P. Ovchinnikov, P. A. Curmi, and D. Pastré, “Role of Microtubules in Stress Granule Assembly: MICROTUBULE DYNAMICAL INSTABILITY FAVORS THE FORMATION OF MICROMETRIC STRESS GRANULES IN CELLS,” Journal of Biological Chemistry, vol. 284, pp. 36569–36580, Dec. 2009.

[25] P. A. Ivanov, E. M. Chudinova, and E. S. Nadezhdina, “Disruption of microtubules inhibits cytoplasmic ribonucleoprotein stress granule formation,” Experimental Cell Research, vol. 290, pp. 227–233, Nov. 2003.

[26] S. Kwon, Y. Zhang, and P. Matthias, “The deacetylase HDAC6 is a novel critical component of stress granules involved in the stress response,” Genes & Development, vol. 21, pp. 3381–3394, Dec. 2007.

[27] M. Loschi, C. C. Leishman, N. Berardone, and G. L. Boccaccio, “Dynein and kinesin regulate stress-granule and P-body dynamics,” Journal of Cell Science, vol. 122, pp. 3973–3982, Nov. 2009.

[28] N.-P. Tsai, Y.-C. Tsui, and L.-N. Wei, “Dynein motor contributes to stress granule dynamics in primary neurons,” Neuroscience, vol. 159, pp. 647–656, Mar. 2009.

[29] E. S. Nadezhdina, A. J. Lomakin, A. A. Shpilman, E. M. Chudinova, and P. A. Ivanov, “Microtubules govern stress granule mobility and dynamics,” Biochimica et Biophysica Acta (BBA) - Molecular Cell Research, vol. 1803, pp. 361–371, Mar. 2010.

[30] E. Vidal-Henriquez and D. Zwicker, “Cavitation controls droplet sizes in elastic media,” arXiv:2102.02506 [cond-mat, physics:physics], Feb. 2021.

[31] P. Ronceray, S. Mao, A. Košmrlj, and M. P. Haataja, “Liquid demixing in elastic networks: Cavitation, permeation, or size selection?,” arXiv:2102.02787 [cond-mat, physics:physics], Feb. 2021.

[32] P. W. Oakes, S. Banerjee, M. C. Marchetti, and M. L. Gardel, “Geometry Regulates Traction Stresses in Adherent Cells,” Biophysical Journal, vol. 107, pp. 825–833, Aug. 2014.

[33] M. J. De Brabander, R. M. Van de Veire, F. E. Aerts, M. Borgers, and P. A. Janssen, “The effects of methyl (5-(2-thienylcarbonyl)-1H-benzimidazol-2-yl) carbamate, (R 17934; NSC 238159), a new synthetic antitumoral drug interfering with microtubules, on mammalian cells cultured in vitro,” Cancer Research, vol. 36, pp. 905–916, Mar. 1976.

[34] Y. Lin, “Nanoparticle Assembly and Transport at Liquid-Liquid Interfaces,” Science, vol. 299, pp. 226–229, Jan. 2003.

[35] L. Keal, C. E. Colosqui, R. H. Tromp, and C. Monteux, “Colloidal Particle Adsorption at Water-Water Interfaces with Ultralow Interfacial Tension,” Physical Review Letters, vol. 120, p. 208003, May 2018.

[36] N. Ballard, A. D. Law, and S. A. F. Bon, “Colloidal particles at fluid interfaces: Behaviour of isolated particles,” Soft Matter, vol. 15, no. 6, pp. 1186–1199, 2019.

[37] A. Gennerich and R. D. Vale, “Walking the walk: How kinesin and dynein coordinate their steps,” Current Opinion in Cell Biology, vol. 21, pp. 59–67, Feb. 2009.

[38] M. Ijavi, R. W. Style, L. Emmanouilidis, A. Kumar, S. M. Meier, A. L. Torzynski, F. H. T. Allain, Y. Barral, M. O. Steinmetz, and E. R. Dufresne, “Surface tensiometry of phase separated protein and polymer droplets by the sessile drop method,” Soft Matter, vol. 17, no. 6, pp. 1655–1662, 2021.

[39] J. W. Cahn and J. E. Hilliard, “Free Energy of a Nonuniform System. I. Interfacial Free Energy,” The Journal of Chemical Physics, vol. 28, pp. 258–267, Feb. 1958.

[40] Y. J. Choi and P. D. Anderson, “Cahn-Hilliard modeling of particles suspended in two-phase flows: CAHN-HILLIARD MODELING OF PARTICLES SUSPENDED IN TWO-PHASE FLOWS,” International Journal for Numerical Methods in Fluids, vol. 69, pp. 995–1015, June 2012.

[41] M. E. Cates and E. Tjhung, “Theories of Binary Fluid Mixtures: From Phase-Separation Kinetics to Active Emulsions,” Journal of Fluid Mechanics, vol. 836, p. P1, Feb. 2018.

[42] Y. Song, U. Shimanovich, T. C. T. Michaels, Q. Ma, J. Li, T. P. J. Knowles, and H. C. Shum, “Fabrication of fibrillosomes from droplets stabilized by protein nanofibrils at all-aqueous interfaces,” Nature Communications, vol. 7, p. 12934, Dec. 2016.

[43] P. M. McCall, S. Srivastava, S. L. Perry, D. R. Kovar, M. L. Gardel, and M. V. Tirrell, “Partitioning and Enhanced Self-Assembly of Actin in Polypeptide Coacervates,” Biophysical Journal, vol. 114, pp. 1636–1645, Apr. 2018.

[44] M. Kothari and T. Cohen, “Effect of elasticity on phase separation in heterogeneous systems,” Journal of the Mechanics and Physics of Solids, vol. 145, p. 104153, Dec. 2020.

[45] Y. Dang, N. Kedersha, W.-K. Low, D. Romo, M. Gorospe, R. Kaufman, P. Anderson, and J. O. Liu, “Eukaryotic Initiation Factor 2*α*-independent Pathway of Stress Granule Induction by the Natural Product Pateamine A,” Journal of Biological Chemistry, vol. 281, pp. 32870–32878, Oct. 2006.

[46] K. E. Kasza, A. C. Rowat, J. Liu, T. E. Angelini, C. P. Brangwynne, G. H. Koenderink, and D. A. Weitz, “The cell as a material,” Current Opinion in Cell Biology, vol. 19, pp. 101–107, Feb. 2007.

[47] G. Guigas, C. Kalla, and M. Weiss, “Probing the Nanoscale Viscoelasticity of Intracellular Fluids in Living Cells,” Biophysical Journal, vol. 93, pp. 316–323, July 2007.

[48] Y.-C. Lin, G. H. Koenderink, F. C. MacKintosh, and D. A. Weitz, “Viscoelastic Properties of Microtubule Networks,” Macromolecules, vol. 40, pp. 7714–7720, Oct. 2007.

[49] H. B. Eral, J. de Ruiter, R. de Ruiter, J. M. Oh, C. Semprebon, M. Brinkmann, and F. Mugele, “Drops on functional fibers: From barrels to clamshells and back,” Soft Matter, vol. 7, no. 11, p. 5138, 2011.

[50] J. Bico, É. Reyssat, and B. Roman, “Elastocapillarity: When Surface Tension Deforms Elastic Solids,” Annual Review of Fluid Mechanics, vol. 50, pp. 629–659, Jan. 2018.

[51] C. Duprat, S. Protière, A. Y. Beebe, and H. A. Stone, “Wetting of flexible fibre arrays,” Nature, vol. 482, pp. 510–513, Feb. 2012.

[52] M. Soleimani, R. J. Hill, and T. G. M. van de Ven, “Capillary Force between Flexible Filaments,” Langmuir, vol. 31, pp. 8328–8334, Aug. 2015.

[53] H. Elettro, S. Neukirch, F. Vollrath, and A. Antkowiak, “In-drop capillary spooling of spider capture thread inspires hybrid fibers with mixed solid-liquid mechanical properties,” Proceedings of the National Academy of Sciences, vol. 113, pp. 6143–6147, May 2016.

[54] M. R. King and S. Petry, “Phase separation of TPX2 enhances and spatially coordinates microtubule nucleation,” Nature Communications, vol. 11, p. 270, Dec. 2020.

[55] S. U. Setru, B. Gouveia, R. Alfaro-Aco, J. W. Shaevitz, H. A. Stone, and S. Petry, “A hydrodynamic instability drives protein droplet formation on microtubules to nucleate branches,” Nature Physics, Jan. 2021.

[56] J. E. Lee, P. I. Cathey, H. Wu, R. Parker, and G. K. Voeltz, “Endoplasmic reticulum contact sites regulate the dynamics of membraneless organelles,” Science, vol. 367, p. eaay7108, Jan. 2020.

[57] J. Agudo-Canalejo, S. W. Schultz, H. Chino, S. M. Migliano, C. Saito, I. Koyama-Honda, H. Stenmark, A. Brech, A. I. May, N. Mizushima, and R. L. Knorr, “Wetting regulates autophagy of phase-separated compartments and the cytosol,” Nature, vol. 591, pp. 142–146, Mar. 2021.

[58] P. Ziltener, A. A. Rebane, M. Graham, A. M. Ernst, and J. E. Rothman, “The golgin family exhibits a propensity to form condensates in living cells,” FEBS Letters, vol. 594, pp. 3086–3094, Oct. 2020.

[59] D. Milovanovic, Y. Wu, X. Bian, and P. De Camilli, “A liquid phase of synapsin and lipid vesicles,” Science, vol. 361, pp. 604–607, Aug. 2018.

[60] S. Elbaum-Garfinkle, “Matter over mind: Liquid phase separation and neurodegeneration,” Journal of Biological Chemistry, vol. 294, pp. 7160–7168, May 2019.

[61] F. G. Quiroz, V. F. Fiore, J. Levorse, L. Polak, E. Wong, H. A. Pasolli, and E. Fuchs, “Liquidliquid phase separation drives skin barrier formation,” Science, vol. 367, p. eaax9554, Mar. 2020.

[62] A. Rai and L. Pelkmans, “Liquid droplets in the skin,” Science, vol. 367, pp. 1193–1194, Mar. 2020.

[63] D. Berchtold, N. Battich, and L. Pelkmans, “A Systems-Level Study Reveals Regulators of Membrane-less Organelles in Human Cells,” Molecular Cell, vol. 72, pp. 1035–1049.e5, Dec. 2018.

[64] G. Cox and C. J. Sheppard, “Practical limits of resolution in confocal and non-linear microscopy,” Microscopy Research and Technique, vol. 63, pp. 18–22, Jan. 2004.

[65] Y. Liu, R. Lipowsky, and R. Dimova, “Concentration Dependence of the Interfacial Tension for Aqueous Two-Phase Polymer Solutions of Dextran and Polyethylene Glycol,” Langmuir, vol. 28, pp. 3831–3839, Feb. 2012.

[66] L. Emmanouilidis, L. Esteban-Hofer, F. F. Damberger, T. de Vries, C. K. X. Nguyen, L. F. Ibáñez, S. Mergenthal, E. Klotzsch, M. Yulikov, G. Jeschke, and F. H.-T. Allain, “NMR and EPR reveal a compaction of the RNA-binding protein FUS upon droplet formation,” Nature Chemical Biology, vol. 17, pp. 608–614, May 2021.

